# Unlocking the functional potential of polyploid yeasts

**DOI:** 10.1101/2021.10.21.465299

**Authors:** Simone Mozzachiodi, Kristoffer Krogerus, Brian Gibson, Alain Nicolas, Gianni Liti

**Affiliations:** Université Côte d’Azur, CNRS, INSERM, IRCAN, Nice, France; Meiogenix, 38, rue Servan, Paris 75011, France; VTT Technical Research Centre of Finland Ltd., Espoo, Finland; Institut Curie, Centre de Recherche, CNRS-UMR3244, PSL Research University, Paris 75005, France

## Abstract

Breeding and domestication have generated widely exploited crops, animals and microbes. However, many *Saccharomyces cerevisiae* industrial strains have complex polyploid genomes and are sterile, preventing genetic improvement strategies based on breeding. Here, we present a novel strain improvement approach based on the budding yeasts’ property to promote genetic recombination when meiosis is interrupted and cells return-to-mitotic-growth (RTG). We demonstrated that two unrelated sterile industrial strains with complex triploid and tetraploid genomes were RTG-competent and developed a visual screening for easy and high-throughput identification of recombined RTG clones based on colony phenotypes. Sequencing of the evolved clones revealed unprecedented levels of RTG-induced genome-wide recombination. We generated and extensively phenotyped a RTG library and identified clones with superior biotechnological traits. Thus, we propose the RTG-framework as a fully non-GMO workflow to rapidly improve industrial yeasts that can be easily brought to the market.

## Introduction

Humans have selected and improved organisms for centuries, leaving profound hallmarks of domestication in their genomes and lifestyles^1^. Selective breeding, in which hybrids with improved performance are generated, was one of humanity’s first biotechnology advances and it is still widely applied^2^. The advent of new genetic engineering techniques enabled to directly manipulate core biological traits relevant for human activities^3^. However, the exploitation of genetically modified organisms (GMOs) in the food sector remains controversial, highly regulated and restricted in many countries.

Humans unwittingly domesticated yeasts of the genus *Saccharomyces* since the earliest food and beverages fermentations were carried out^4–6^. Since then, novel fermentation processes selected new yeast strains that are still used nowadays^7^. Nevertheless, there is a pressing interest across different sectors that exploit *Saccharomyces* yeasts to create new strain variants able to better tolerate stresses encountered during the industrial fermentations or increase the yield of desired compounds. Newly designed *de novo* lab-hybrids that combine different *Saccharomyces* species have displayed good or superior fermenting properties compared to common commercial strains suggesting that indeed commercial strains are not yet optimized, leaving room for improvement despite the long domestication period^8–10^. However, attempts to cross industrial yeasts through selective breeding or induce genomic recombination through meiosis are often unsuccessful due to the inherent sterility of the strains, which is a hallmark of yeast domestication^6,11^. This domestication-syndrome might have derived from the lack of selection on sexual reproduction, random genetic drift or adaptation to specific fermentation niches, leading to the accumulation of punctuated deleterious loss-of-functions (LOF) alleles that inactivate genes involved in the gametogenesis (sporulation in yeast). In addition, the genomes of industrial strains often have features such as extreme sequence divergence between the subgenomes (namely heterozygosity), structural rearrangements, aneuploidy and polyploidy all of which are known to contribute to sterility^12^. Classic examples are strains used in beer production or in the baking industry sharing different ancestries that have repeatedly converged toward polyploid complex genomes^5,13,14^. However, tetraploid strains derived from designed crosses perform a correct chromosome segregation and produce viable gametes^15^ in contrast to triploid strains. Therefore, the genetic basis driving this extreme sterility are not yet fully understood and multiple factors likely contribute to impair the sexual reproduction of industrial polyploid strains. Thus, other approaches to improve sterile industrial strains have been proposed^16^ because directly fixing the lack of a complete sexual reproduction remains unfeasible.

Recently, we demonstrated that aborting meiosis in laboratory-derived sterile diploid hybrids and returning them to mitotic growth, a process called return-to-growth (RTG), generated genetic diversity and resulted in phenotypic variability in the evolved samples^17,18^. RTG is induced when cells that have entered meiosis but are not yet committed to complete it are shifted back to a nutrient-rich environment^19^. As in normal meiosis, after DNA replication Spo11p induces multiple genome-wide double-stranded breaks (DSBs) that lead to the formation of intermediate recombinant molecules^20^. These molecules are then resolved upon the resumption of the mitotic cycle, resulting in dispersed LOH tracts that derive from the segregation of the chromatids in the mother and the daughter cells. Despite generating multiple dispersed LOH tracts, the RTG process is not mutagenic and it preserves the initial genome content in diploid laboratory strains^17^. Furthermore, the LOHs induced by RTG can lead to rapid phenotypic diversification through the unmasking of beneficial alleles^17,18^. Given that RTG induces phenotypic variation without complete sexual reproduction, which often is defective in industrial strains, RTG may represent a powerful approach for improvement of industrial strains. However, it is still unknown whether the RTG paradigm can induce recombination and unlocks novel phenotypic variability without triggering systemic genomic instability in such complex genomic scenarios. Furthermore, the selection of RTG recombinant clone either requires genetically-engineered selective markers^17^ or a lengthy and low throughput microdissection approach^18^. This hinders the scaling-up of the RTG process to generate a large library of recombined industrial yeasts. Here, we developed a comprehensive workflow to improve polyploid industrial yeasts through RTG and easily select RTG-recombined clones. We applied this workflow to generate a non-GMO library of evolved industrial polyploid strains harbouring improved biotechnological traits.

## Results

### Genomic and reproductive portraits of industrial polyploid yeasts

Domesticated strains derived from wild yeast ancestors show hallmarks of genomic complexity such as polyploidy, aneuploidy and horizontal gene transfer (HGT) (**Fig. 1a**). In order to develop a tailored RTG framework for industrial strains (**Fig. 1b**), we selected two genetically unrelated industrial *S. cerevisiae* strains as test cases, hereafter called OS1364 and OS1431 (**Fig. 1a**), and characterise their genomes and reproductive capacity. OS1364 was isolated from a cassava factory in Brazil and belongs to the mosaic beer clade, whereas OS1431 was isolated from an ale beer fermentation in England and belongs to the ale beer clade^13^ (**Supplementary table 1**). We performed both short- and long-reads sequencing and detected multiple hallmarks of domestication. Despite their genetic ancestry, both strains are polyploid, with OS1364 being triploid (3n, with one additional copy of chromosome III) and OS1431 being tetraploid (4n) (**Supplementary Figure 1a-b**). Furthermore, both genomes harbour a considerable number of heterozygous positions (~40K) distributed genome-wide suggesting that these two strains are intraspecies hybrids generated by admixture of different *S. cerevisiae* strains (**Fig. 1c, Supplementary Figure 1c**). A large region of loss-of-heterozygosity (LOH) is present on chromosome XII downstream of the rDNA locus in both strains (**Supplementary Figure 1c),** consistent with this locus being inherently prone to recombination^5,13^. We detected several single-nucleotide polymorphisms within coding regions (*n*=28932 OS1364, *n*=32399 OS1431) with OS1431 harbouring more than the expected number (Expected ~75%, based on the reference genome, Observed 82.2%). Among these polymorphisms we found pervasive heterozygous missense variants (*n*=10747 OS1364, *n*=13033 OS1431, **Supplementary Figure 1d**). We observed that the genetic variation also manifests in the form of copy number variants (CNVs) (**Supplementary Figure 1b**), resulting in small amplifications or deletions similar to those previously observed in industrial strains^5^. Subtelomeric regions are known to be highly enriched in this type of variation^21^. For instance, we detected a large horizontal gene transfer (HGT) region close to the subtelomere XIII-R of OS1364 (present in two homologs) (**Supplementary Figure 1e**). HGT regions are known to harbour genes that improve fitness in the fermentation environment and have been detected often in domestic strains^22^. The genetic diversity observed in the two strains might reflect the different evolutionary histories of the haplotypes before or after the admixture.

**Fig. 1.**
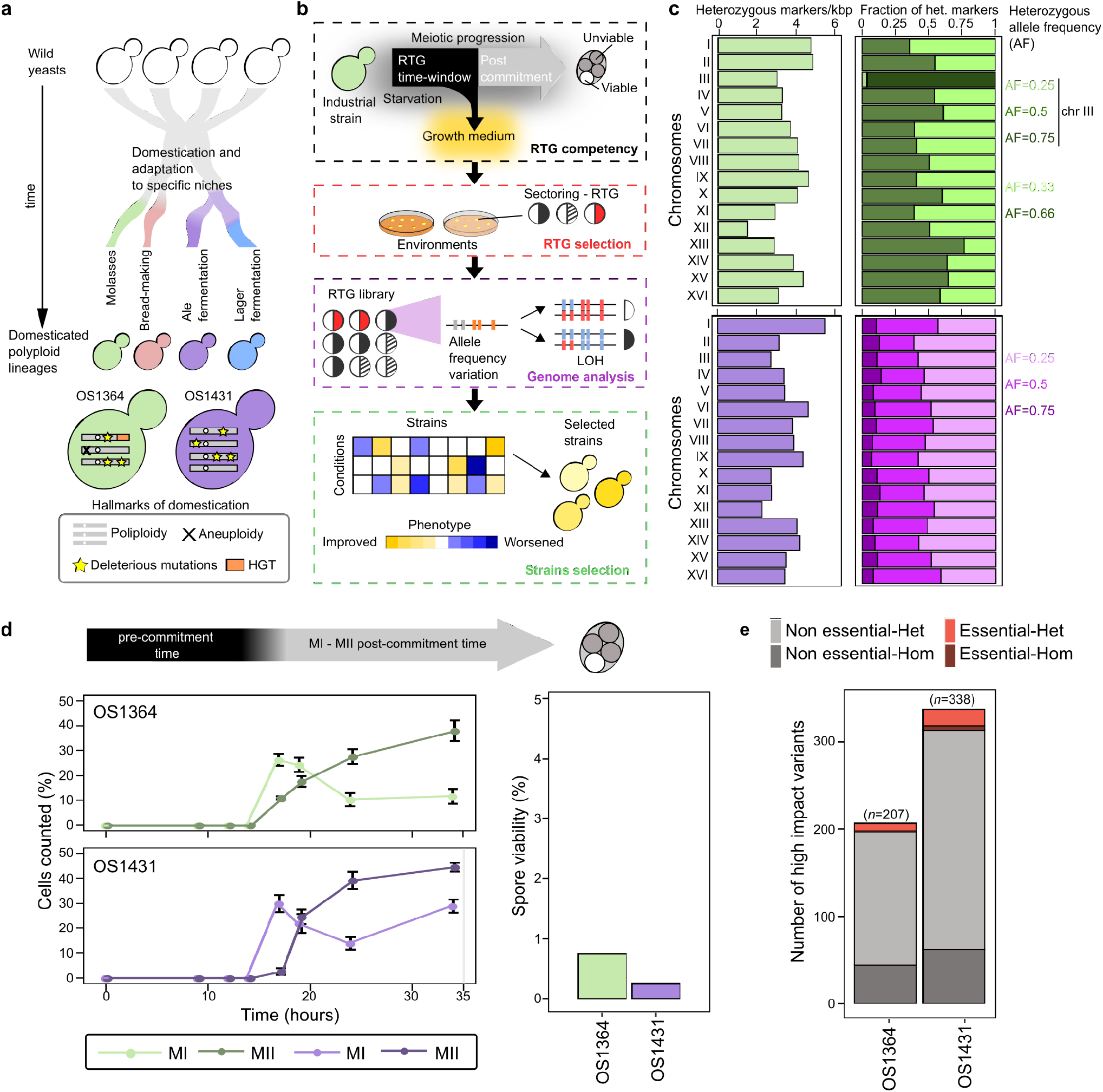
Genome complexity and sterility of domesticated *S. cerevisiae*. **(a)** Domestication produced polyploid strains adapted to specific human-made niches that are characterised by common genomic hallmarks (bottom box), which are detected in both strains used in this study. **(b)** Schematic depicting the RTG framework: we initially quantify and optimise the meiotic progression, followed by selection of novel colony phenotypes. Finally, RTG samples are sequenced and phenotyped to identify clones with improved industrial traits. (**c**) Left: Level of heterozygosity (number of heterozygous markers/kbp) across each chromosome, total number of heterozygous markers (*n*=40431 OS1364 (green), *n*=39376 OS1431 (purple)). Right: Heterozygous markers partitioned according to their allele frequency. (**d)** Meiotic progression measured by nuclei DAPI staining, reported as the average (*n*=3 replicates) of the percentage of cells that have progressed over the first (MI) and second (MII) meiotic division. The error bars represent the standard deviation. On the right, bar plot representing the spore viability (3 viable spores/400 OS1364, 1 viable spore/400 OS1431). The top arrow is a qualitative description of the meiotic progression based on the results of the DAPI staining. (**e**) The bar plot represents the number of high-impact variants affecting essential (red) or non-essential (grey) genes and further divided into homozygous (Hom) if present in all the haplotypes, and heterozygous (Het) if not present in all the haplotypes.

Next, we investigated the reproductive capacity of these polyploid strains and observed a defective sexual reproduction (**Fig. 1d**). Specifically, the strains showed slow, asynchronous and inefficient meiotic progression and generated nearly completely unviable gametes (less than 1%). The high heterozygosity and lack of an effective sexual reproduction suggest that the genetic load might contribute to gametes’ unviability. By using Ensamble Variant Effect Predictor (VEP) (**Methods**), we detected many highly deleterious variants in both genomes (start-loss, stop-loss, stop-gain, *n*=207 OS1364, *n=338* OS1431), including 10 (OS1364) and 24 (OS1431) in essential genes^23^ (**Fig. 1e**, **Supplementary Tables 2 and 3**). OS1364 has accumulated a number of deleterious variants not significantly different from the average number of LOFs observed in domesticated strains of the 1011 *S. cerevisiae* collection^11^, whereas OS1431 has accumulated significantly more LOFs (**Supplementary Figure 1f**). Given OS1431 is tetraploid and tetraploid strains correctly segregate their chromosomes in contrast with triploid strains^24^, the observed genetic load is likely to drive part of its sterility. However, we performed a GO-term analysis to test if genes involved in the sporulation process had accumulated LOFs and we did not find any enrichment of LOF in sporulation associated genes in frame with the results that cells of both strains entered meiosis when starved. Overall, these two sterile domesticated *S. cerevisiae* strains with complex polyploid genomes represent ideal test cases to probe the RTG framework.

### Sterile polyploids are RTG competent

Despite the fact these polyploid *S. cerevisiae* hybrids show an inefficient meiotic progression and generate inviable gametes, we conjectured that RTG should not be precluded since it only relies on the meiotic prophase progression, a time window in which cells are not yet committed to complete meiosis. To test if our two polyploid hybrids are able to perform RTG, we engineered them with a genetic system to measure LOH rates at a heteroallelic *LYS2/URA3* locus on chromosome II, that we have broadly applied in RTG experiments^17,25^ (**Fig. 2a**). We validated the genotype of the engineered *LYS2/URA3* strains by PCR and growth on selective media (**Supplementary Figure 2a, Supplementary Table 4**). Then, we evolved the engineered OS1364^*LYS2/URA3*^ and OS1431^*LYS2/URA3*^ through RTG and collected cell populations throughout the meiotic progression (**Fig. 2b**). We calculated the RTG-induced recombination by comparing the basal level of cells growing on 5-FOA measured in the unsporulated cultures (T0) to the RTG cells obtained after 6 hours (T6) and 14 hours (T14) of sporulation induction (**Methods**). We detected a 10-fold and 3-fold increase of cells growing on 5-FOA, indicating an increased LOH rate at T14 in OS1364^*LYS2/URA3*^ and in OS1431^*LYS2/URA3*^ respectively, whereas the increase was not significant at T6, consistent with the fact that cells have not progressed sufficiently through meiosis at this timepoint (**Fig. 2b, Supplementary Table 5**). The absolute percentage of cells grown at T14, calculated subtracting the T0 background value to the T14 (**Methods**), was almost 2-fold higher in OS1431^*LYS2/URA3*^ (0.10 % in OS1364^*LYS2/URA3*^, 0.19 % OS1431^*LYS2/URA3*^), in line with its lower heterozygosity on chromosome II (**Fig. 1c**) where pre-existing small LOH regions might represent preferential sites of recombination^25^. To further prove that RTG induced the increase in recombination observed at the *LYS2/URA3* locus we deleted all three copies of *SPO11*, which is essential for inducing DSBs in meiosis, in OS1364^*LYS2/URA3*^ (**Supplementary Figure 2b)** and measured recombination. We did not detect any significant increase between the T0 and the respective T14 in the OS1364^*LYS2/URA3*^ *spo11Δ* strain (**Supplementary Figure 2c, Supplementary Table 5**), supporting the conjecture that RTG caused the increased recombination as it relies on the Spo11p induced DSBs in early meiosis.

**Fig. 2.**
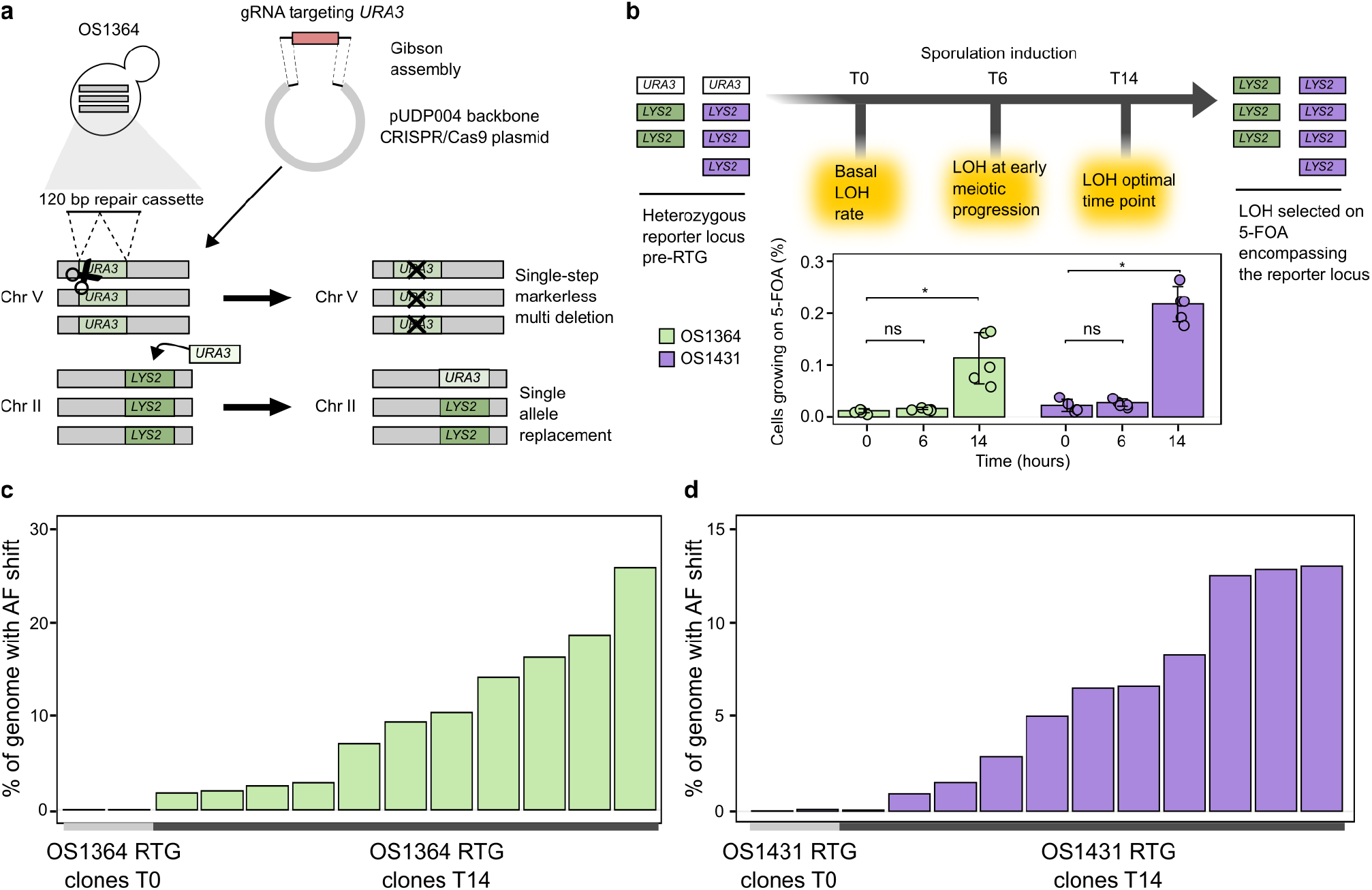
RTG induced recombination in polyploid sterile hybrids. **(a)** A CRISPR-based approach enabled the engineering of the *URA3-*loss genetic system in the polyploid strains. The strains generated in such way are hereafter referred to as *LYS2/URA3* regardless of the number of loci. **(b)** Reporter assay used to measure recombination rate at the heterozygotic locus *LYS2/URA3* (top panel). Bar-plots representing the average (*n*=5 replicates) percentage of cells growing on 5-FOA at different time points: no sporulation induction (T0), 6 hours (T6) and 14 hours (T14) of sporulation induction (bottom panel). The increase of cells growing on 5-FOA at T14 compared to T0 was significant for both samples (*p*-value < 0.05, one-sided Wilcoxon ranked-sum test) whereas was not significant at T6 (*p*-value > 0.05, one-sided Wilcoxon ranked-sum test). The error-bars represent the standard deviation. (**c-d**) Bar plot representing the percentage of genome in which we detected an allele frequency (AF) shift for the RTG samples derived from OS1364^*LYS2/URA3*^ (green, **c**) and OS1431^*LYS2/URA3*^ (purple, **d**). Recombination and LOH induced upon genome engineering with CRISPR/Cas9 at the native *URA3* locus was excluded.

We next performed whole-genome sequencing of the *LYS2/URA3* parental strains, controls (T0) (*n=2* OS1364, *n*=2 OS1431, **Table S1)** and RTG-evolved clones (T14) (*n*=11 OS1364, *n*=11 OS1431, **Table S1**) isolated on 5-FOA plates to evaluate the genome-wide impact of RTG. Our analyses revealed varying levels of recombination in the RTG clones, consistent with previous reports in RTG diploid hybrids^17,18^ (**Fig. 2c-d, Supplementary Figure 2d**). The RTG clones derived from OS1364^*LYS2/URA3*^ had an average of 10% (maximal value 26%) of the genome affected by LOH, whereas OS1431^*LYS2/URA3*^ had an average of 6.7% (maximal value 13.5%). Moreover, we did not detect any chromosome loss potentially accounting for 5-FOA resistance in the RTG sequenced clones, showing that despite the strain polyploidy, LOH at the *LYS2/URA3* locus arose more frequently than aneuploidy during RTG. We observed overall genome-wide stability and detected only one aneuploidy and two large CNVs across the 11 RTGs derived from OS1431^*LYS2/URA3*^, where the two CNVs likely resulted from ectopic recombination initiated by a small region of homology between chromosome IX and chromosome XIV (**Supplementary Figure 3a-b-c**). Similarly, we detected only two aneuploidies in one RTG sample across the 11 samples derived from OS1364^*LYS2/URA3*^. Altogether, these results demonstrate that these polyploid hybrids are RTG-competent despite their meiotic sterility, supporting that the RTG workflow can generate genetic diversity in sterile industrial strains.

### Selection of recombined RTG clones by natural colony phenotypes

The RTG selection based on *URA3-loss* requires genome editing and produces GMOs that have marketing constraints in the food industry. To overcome this limitation, we devised an alternative selection strategy based on natural variability in colony phenotypes such as morphology and colour. Colony phenotypes are complex traits, influenced by both genetic and environmental factors (**Fig. 3a)**. We used a YPD-based medium with varying concentrations of dextrose (0.5% and 1%), a major environmental regulator of colony morphology^26,27^, to unveil novel phenotype of colonies derived from RTGs of OS1364 and OS1431 wild-type (WT) cells (**Fig. 3a, Methods**). Indeed, we found phenotypic variability in the RTG colonies but the divergent phenotype was often present in only a large sector (approximately half) of the colony. Therefore, we hypothesized that the sectored colonies were mother-daughter (M-D) RTG pairs that did not complete budding at the time of plating. Consistently, we found sectored and non-sectored colonies according to when cells were plated after the RTG induction (**Supplementary Figure 4a**). We observed three types of sectored colony phenotypes in the RTG plates of OS1364 (wrinkled-smooth, red-white and dark-white) (**Fig. 3b, Supplementary Figure 4, Supplementary Table 5**) with a summed total frequency of ~0.75 % and one type of sectored colony in the OS1431 RTGs (wrinkled-smooth) with a frequency of ~0.5 % across all environments tested (**Fig. 3b, Supplementary Table 5**). In contrast, we did not observe any of these phenotypes in control plates where cells were plated without inducing sporulation. Thus, this result was in line with our initial hypothesis that the sectored colonies were unveiled in the modified YPD plates upon budding of recombined mother-daughter (M-D) RTGs, whereas the non-sectored colonies displaying phenotype variation represent RTG cells in which the mother and daughter were separated by the first budding before the plating. An alternative explanation is that the sectored RTG colonies derived from residual sporulation and spore germination, although this is unlikely given the near to zero spore viability observed. To undoubtedly exclude this scenario, we deleted the *NDT80* gene by CRISPR/Cas9 multi-deletion in both hybrids, to generate mutants that are RTG competent^17,20^ but arrest before the first meiotic division (MI) and therefore unable to complete sporulation (**Supplementary Figure 5a-b**). Then, we evolved the OS1364^*ndt80Δ*^ and OS1431^*ndt80Δ*^ mutants through the same RTG protocol used for the wild-type hybrids (**Supplementary Figure 5b**) and detected sectored phenotypes similar to those seen in the WT hybrids, thus excluding the possibility that residual sporulation contributes to the formation of sectored colonies.

**Fig. 3.**
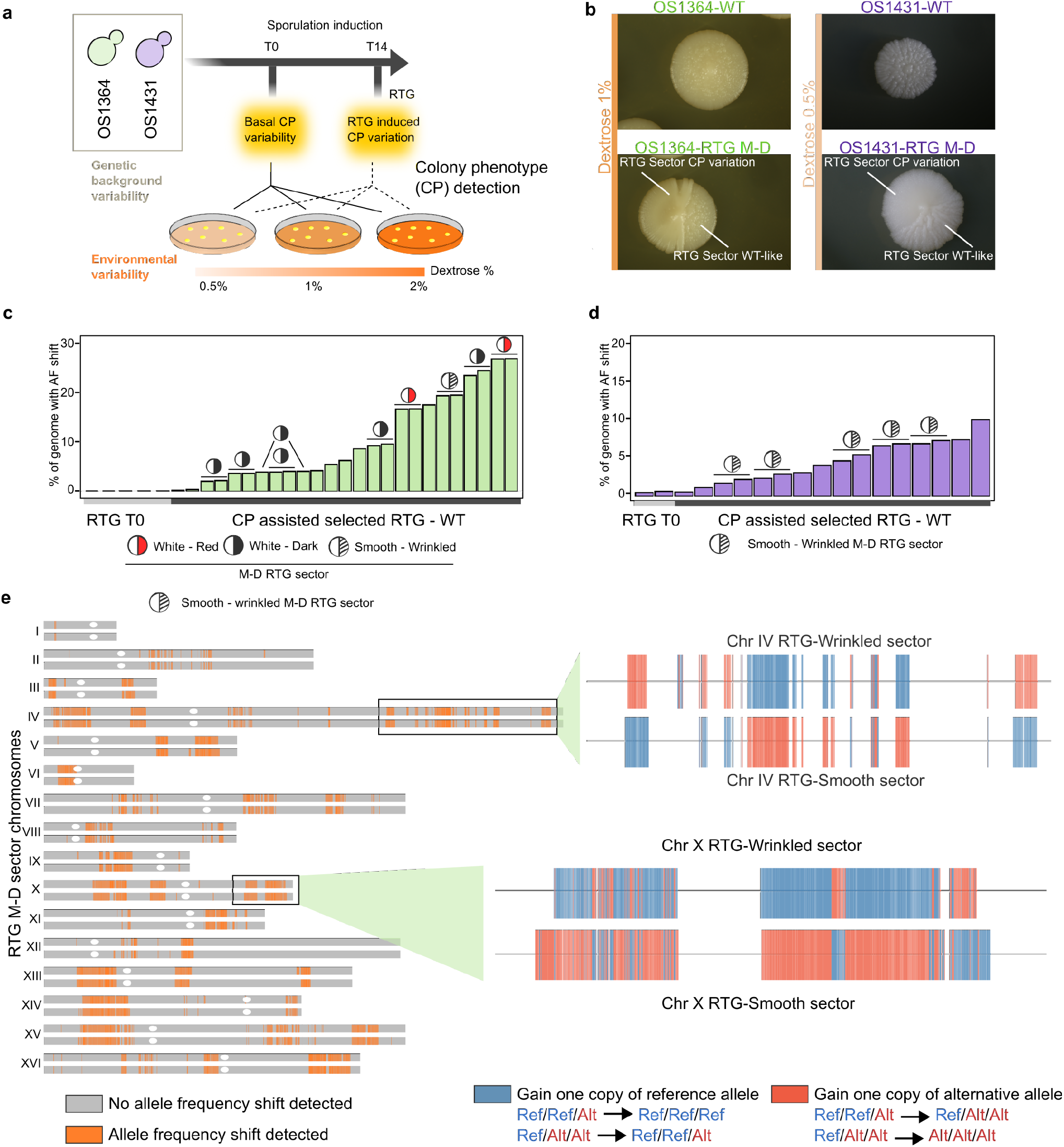
Colony phenotype variability revealed mother-daughter RTG pairs. **(a)** Varying dextrose concentrations unveiled hidden colony phenotypic variation upon RTG. **(b)** Wild-type (WT) colony phenotypes (top) and sectored phenotypes emerging on RTG plates (bottom). The concentration of dextrose in the media is reported on the left. **(c)** The percentage of markers with allele frequency (AF) shift in the OS1364 derived RTG samples. The mother-daughter pairs for which both sectors were sequenced are indicated (circles). Single samples represent non-sectored RTG colonies (*n*=4) or only one-side of a RTG sectored colony (*n*=3). **(d)** As in panel **c** for the OS1431 background. **(e)** Recombination map of a RTG M-D pair (top chromosome: sm244/ bottom chromosome: sm245, derived from OS1364). Grey regions indicate the heterozygous markers without AF shifts, whereas orange regions contain heterozygous markers with AF shifts underlying LOH events. A zoom-in of two recombined regions is illustrated on the right. The colour code represents the genotype variation of the heterozygous markers so that the gain of a reference allele given the initial genotype is indicated in blue, whereas the gain of an alternative allele is indicated in red.

We sequenced multiple paired samples derived from sectored RTG colonies and non-sectored RTG colonies derived from RTG in the OS1364 and OS1341wild-type and *ndt80Δ* strains and samples isolated from control plates (T0) (**Supplementary Table 1**). Whole-genome sequencing revealed recombination detected as allele frequency (AF) shifts across the unphased heterozygous markers in all the putative M-D RTG colonies with a sectored phenotype (**Fig. 3c-d, Supplementary Figure 5c-d**). The fraction of the genome in which we detected RTG-induced recombination was highly variable in both strains (9.7±8.2% in OS1364, 4.8±2.4% in OS1431 WT-RTG, average ± SD), mirroring the results obtained from the *URA3*-loss assay. Moreover, we provided two additional proofs that the sectored RTG colonies were genuine RTG M-D pairs. First, the AF shift for heterozygous markers was largely reciprocal in the wild-type and *ndt80Δ* sectored RTG pairs with the rest of the non-reciprocal events representing gene-conversion (**Fig. 3e, Supplementary Figure 6a-b-c, Supplementary Table 12**) as found in M-D RTG pairs of diploid lab strains. In addition, by comparing each pair-wise combination of RTG M-D pairs we also found that each pair is clearly differentiated from the others in term of recombination landscape (**Supplementary Figure 6d**).

Nevertheless, some of these regions of recombination were shared across RTG M-D pairs with the same sectored phenotype (**Supplementary Figure 7a**) supporting the scenario that the colony phenotypes observed were driven by localised LOH. Second, we observed a complementary gain and loss of one chromosome in one M-D pair derived from OS1364 (**Supplementary Figure 7b**), which is compatible with chromosome missegregation between M-D during RTG. Thus, we concluded that the sectored RTG colonies were indeed M-D pairs and we exploited the recombination generated by RTG in OS1364 M-D RTGs to produce phased local haplotypes showing a potential application of RTG in polyploid yeasts (**Supplementary Figure 7c**). Overall, we showed that highly recombined RTG M-D pairs can be selected by exploiting natural colony phenotypic variation and we proved that these phenotypes arise as a result of RTG-induced recombination.

### RTG recombination improves industrial fitness

Next, we probed the potential of RTG to improve industrial phenotypes. First, we compared the fermentation performances of OS1364 and OS1431 to the commercial strain WLP001 used as a reference to evaluate the fermentation performances of our strains (**Supplementary Table 1**). We confirmed that our two selected strains are competitive for industrial fermentation, with OS1364 outperforming both WLP001 and OS1431 for fast fermentation (**Supplementary Table 10**). However, both OS1364 and OS1431 had a substantial decline in cell viability (of ~25% and ~50% respectively) often observed in chronologically aged polyploid *S. cerevisiae^11^*, which is more modest in WLP001 (8%, **Supplementary Table 11**).

We started our phenotypic screening to identify potential improved RTG variants by evaluating selected non-GMO RTGs (*n*=25 OS1364, *n*=16 OS1431, **Supplementary Table 6**) and compared them to their respective parental strain and T0 controls (*n*=2 OS1364, *n*=2 OS1431) across several conditions mirroring industrial fermentations (**Fig. 4a**). We performed a first phenotypic screening in osmotic and alcoholic stress conditions and found that RTG samples had broad phenotypic variability with either a worsened, unchanged or improved phenotype as compared to the parental strains and the T0 controls (**Fig. 4b, Supplementary Table 6**). This scenario is consistent with the LOHs induced by RTG arising randomly in the genome and not as a by-product of a specific selective pressure. Moreover, some M-D RTG pairs showed complementary growth-rate variation (**Fig. 4c**) that can be explained by complementary LOHs segregating weaker and stronger alleles in the RTG pair. Growth rate variation in ethanol and maltose appeared to be moderately correlated in the OS1364 RTG library, underscoring that recombination might have affected pleiotropic genes regulating both traits (**Supplementary Figures 8a-b**).

**Fig. 4.**
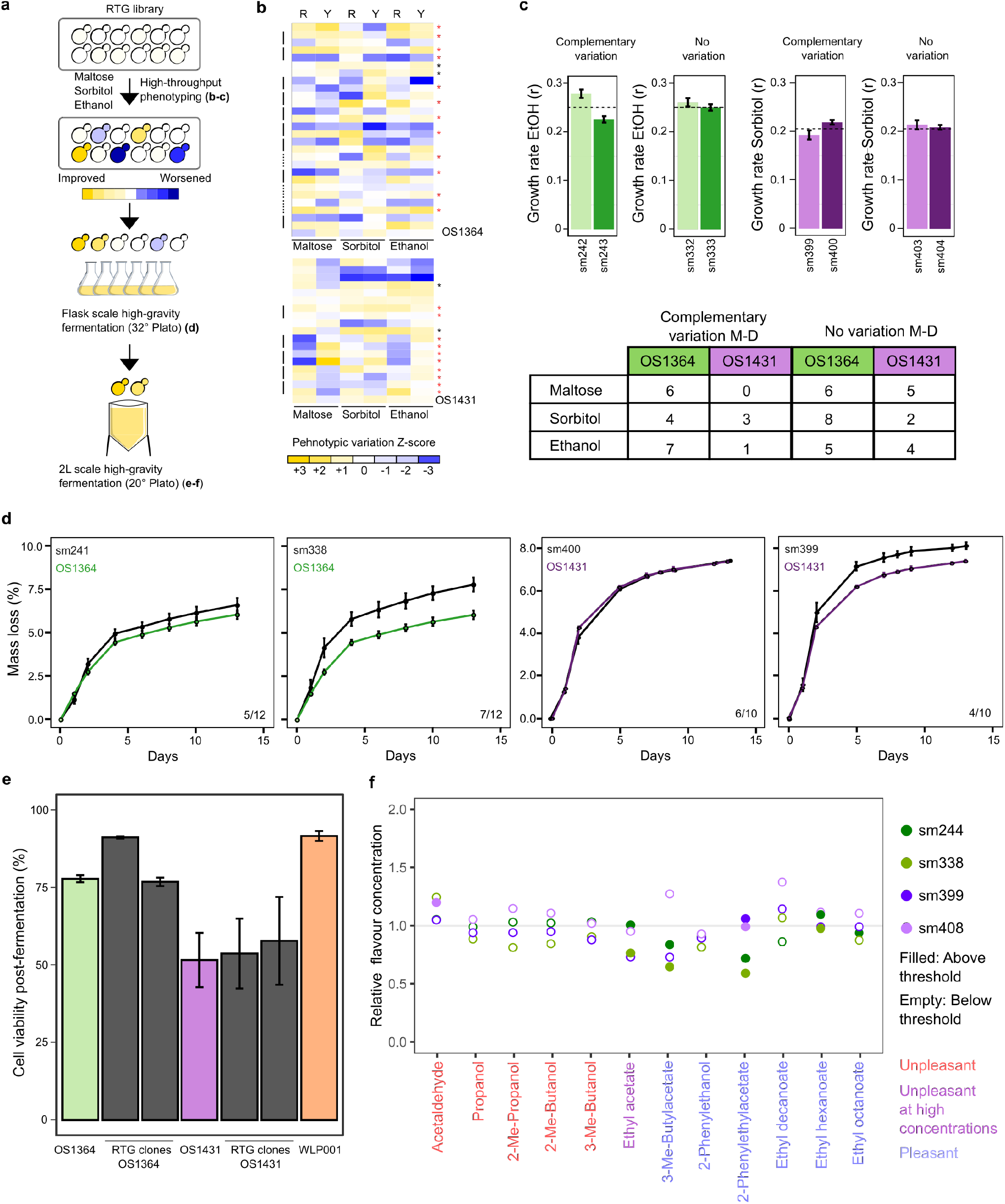
Phenotypic variability in industrial RTG variants. **(a)** Schematic of the phenotypic screening performed. Letters in parentheses indicate the respective data for the phenotypic screening depicted **(b)** Heatmaps representing growth rate (1^st^ column, R) and maximal optical density (2^nd^ column, Y) of the RTG samples selected for the next screening (red star), T0 samples (black star) and the M-D sectors (connected with lateral black line) and RTG-sectors for which only one sample of the sector (dotted-black line). Each measure is the average of 4 technical replicates. **(c)** Phenotypic diversification in sectored M-D pairs of OS1364 (green) and OS1431 (purple). Bar plots represent an example of an RTG M-D pair with divergent phenotypic variation and absence of variation. A similar example is reported on the left for OS1431. The dotted line represents the average phenotype in the RTG M-D library and the error bar represents the standard deviation. On the bottom table is reported the number of M-D pairs showing a complementary phenotypic variation per each class. **(d)** Mass loss curves during the flask scale fermentation experiment of an RTG without variation (left) and an improved RTG (right) for OS1364 (green) and OS1431 (purple), the error bars represent the standard deviation. The numbers on the bottom right represent the RTG samples in each category on the overall number of RTGs screened. **(e)** Post fermentation viability measured after a 2L-fermentation in high gravity wort for the two parents, the respective RTG samples and a commercial strain in triplicate. One RTG (sm244) derived from OS1364 showed a huge increased in viability. The error bars represent the standard deviation. **(f)** Variability in the aroma profile across the four RTGs from the 2L scale fermentation, the values are expressed as relative change compared to the concentration produced by the parental strain.

Next, we selected RTG samples derived from OS1364 based on their phenotypic performance (*n*=12) and their LOH landscape across a core set of fermentation genes (*n*=74), and all the M-D RTG pairs from OS1431 (*n*=10) (**Supplementary Tables 7 and 8**). We inoculated the selected RTGs in a high-gravity wort (32° Plato), representing extreme fermentative conditions, to further investigate differences in stress resistance, carrying out a flask-scale fermentation for 13 days evaluating mass loss and ethanol produced at regular intervals (**Supplementary Table 9**). Some RTG samples showed increased fermentation kinetics (**Fig. 4d**) as well as a superior alcohol production (**Supplementary Figure 8c**) compared to the parental strains. We also noticed that in some M-D RTGs derived from OS1431 the fermentation performances were complementary (**Supplementary Figure 8d, Supplementary Table 9).** We hypothesized that RTG samples showing improved fermentation performances should also have improved resistance to fermentation stress and, potentially, higher post-fermentation cell viability. Therefore, we selected two RTG variants for each background among the best performers in the previous flask-scale fermentation, to carry on a 2-liter scale fermentation in high-gravity wort, to mimic modern industrial brewery practice (20° Plato). We found that OS1364 RTGs performed at least equally well as their respective parental strains, while OS1431 RTGs were slightly better in fermentation performances (**Supplementary Figure 9a, Supplementary Table 10**) but did not have increased post-fermentation viability (**Fig. 4e**). Remarkably, one OS1364 RTG had a large increase in post-fermentation cell viability **(Fig. 4e, Supplementary Table 11)**, reaching a level similar to WLP001, showing that RTG can fix detrimental traits.

Furthermore, we evaluated the aroma profile and the post-fermentation residual sugars, two parameters that are highly relevant in the beer industry. The two parental strains produced distinct aroma profiles from that of WLP001 and consumed almost all the sugars present in the wort (**Supplementary Table 11**). The four selected RTG samples did not show any deleterious trade-off in either phenotypes with the exception of a slight increase of acetaldehyde, in sm408, which is not desirable but the increase was negligible (Sensory threshold = 10 mg/mL, sm408 = 10.5 mg/mL, **Fig. 4f**, **Supplementary Table 11**). One RTG derived from OS1364 had a lower production of ethyl acetate (**Fig. 4f**, **Supplementary Figure 9c, Supplementary Table 11**). This is consistent with recombination encompassing the gene *ATF2*, which has a role in shaping this trait (**Supplementary Table 8**). Moreover, the RTG derived from OS1364 showed diversified production of esters, which may lead to a further differentiation of the sensory profile of the beer **(Fig. 4f)**.

Overall, our data demonstrate that RTG recombination in sterile polyploid strains can unlock phenotypic improvement in traits of industrial relevance^16^, contributing toward microbial stability or shaping the sensory quality of the product.

## Discussion

In this work, we showed that RTG is an efficient approach to generate genetic and phenotypic diversity in two fundamentally different industrial sterile yeast strains. Current approaches to improve industrial yeasts largely depend on designed targeted genetic modifications. Such approaches must contend with market restrictions and can garner societal mistrust^16^. As an alternative, we now show that RTG-assisted homologous recombination is an efficient approach to generate genetic and phenotypic diversity in polyploid and sterile industrial yeasts. The RTG approach, coupled with selection based on natural phenotypes generates non-GMO yeast strains that can be unrestrictedly introduced into the market. Compared to other non-GMO approaches such as serial transfer^28,29^, RTG induces a random genome shuffling as it does not select for a specific trait except for the genetic loci regulating the natural phenotype selected. Nevertheless, RTG has several benefits over the previous improvement strategies. We showed that the RTG process does not trigger genome instability in polyploid strains, similarly to diploid hybrids^17,18^. This is in contrast with direct selection where the abundant number of generations performed often results in ploidy and chromosome copy number variation^30^. Furthermore, trade-offs in unselected traits have been described as undesirable outcomes in adaptive evolution experiments^31–33^, although some solutions have been proposed to fix, for instance, the impaired sugar consumption emerging after adaptive evolution^34^. We also observed RTG variants with impaired performances. In fact, we found that the RTG M-D pairs often had complementary worsened and improved fitness in different environments. In addition, multiple deleterious trade-offs can accumulate in RTG samples when the fraction of the genome that recombines is large. However, the RTG library harbours samples with variabile levels of genome-wide recombination and samples with a low level of recombination might be less likely to experience undesired trade-offs. Nevertheless, one of the most recombined RTG did not show phenotypic decay in the screenings performed; on the contrary, it was better able to tolerate the harsh fermentation conditions and had increased post-fermentative viability.

The RTG libraries are produced without introducing any selective pressure. Therefore, they are phenotypically agnostic, except for the trait used for the selection and can harbour improved phenotypes that are useful for multiple applications. The variability of “colony-associated-phenotypes” is a key parameter of the approach we devised for selecting M-D RTG recombinants, and it might be limited by the variability of natural phenotypes that can be screened upon RTG induction. However, a simultaneous screening of several environments could provide an effective solution to unveil variability in colony phenotypes. For instance, there are media which can trigger a colour variation linked to relevant industrial traits, such as the production of hydrogen sulfide detected in BiGGY agar plates^35^. Moreover, our approach successfully captured mother and daughter RTG pairs that have only been isolated so far with a micromanipulation approach^18^ developed for laboratory strains that cannot be scaled-up. We envision that our selection method can be easily automated to quickly screen hundreds of colonies^36^. In addition, our selection protocol can be easily performed and seamlessly integrated with other RTG methods that do not feature mother-daughter RTG pairs^37^ or other yeast improvement strategies. For instance, strains derived from an adaptive evolution experiment could be further improved through RTG to erase trade-offs, or, vice-versa, RTG samples could be submitted to adaptive evolution to be further optimized. In addition, the RTG framework can be easily generalized to reshuffle designed hybrid genomes^17,18^ as the only requirement is that the hybrid can enter meiosis and progress until the first meiotic prophase, even if with a low efficiency. The RTG approach has other practical applications as RTG libraries can be used in large-scale linkage analyses to unravel the genetic architecture of complex industrial traits and the RTG induced recombination could aid existing genome phasing methods^38^ in producing polyploid *de-novo* genome-assemblies. In conclusion, we propose the RTG framework as a novel avenue to induce genetic and phenotypic variability in industrial sterile yeasts. RTG can generate variants that can easily reach the market and can also serve as a method to help our understanding of complex genetic traits in industrial strains.

## Methods

### Detection of deleterious, missense mutations and pre-existing LOHs

Short reads of the parental strains were obtained from the 1011 *S. cerevisiae* yeast project^13^ and mapped against the SGD reference R64.2.1 using bwa-mem algorithm. Single nucleotide variants were called by using Freebayes (v1.3.1-19) with the argument “-p” to set an appropriate ploidy and quality >20. We annotated impactful mutations by using the Variant Effect Predictor (VEP) suite^39^. Mutations were annotated as impactful if they caused a stop-loss, stop-gain or start-loss. Subtelomeric variants or frameshift ones were included in the list to make the data comparable to the table of LOFs generated for the 1011 *S. cerevisiae* yeast project^13^. The list of essential genes was obtained from Liu et al^23^. The GO-term analysis was performed using the SGD GO-term analysis suite at https://www.yeastgenome.org/ with a *p*-value threshold of 0.01 and selecting GO term associated with sporulation. We identified pre-existing LOHs in the genome of the parental strains by searching for non-overlapping regions of 50 kbp where we identified 10 or less heterozygous markers that where not shared by all the unphased haplotypes. The plots were generated using ggplot2 and in-house R scripts.

### Genome content analysis

The genome content of OS1364 and OS1431 was measured using propidium iodide staining as described in Mozzachiodi et al^17^ and compared to a reference diploid strain. Each strain was patched from a −80°C glycerol stock onto a YPD plate (1% yeast extract, 2% peptone, 2% dextrose, 2% agar) and incubated overnight at 30 °C. The following days the strains were incubated in 1 mL liquid YPD medium (1% yeast extract, 2% peptone, 2% dextrose) and growth overnight at 30 °C without shaking. The next day 200 μL of the overnight culture was resuspended in 1 mL of fresh YPD and growth until exponential phase. Then cells were centrifugated, washed with 1 mL water and fixed overnight in 1 mL of EtOH 70%. The following day each sample was washed with PBS 1X, resuspended in the PI staining solution (15 μM PI, 100 μg/mL RNase A, 0.1% v/v Triton-X, in PBS) and incubated for 3 hours at 37 °C in the dark. Ten thousand cells for each sample were analysed on a FACS-Calibur flow cytometer. Cells were excited at 488 nm and fluorescence was collected with a FL2-A filter. An increase of 50% of the fluorescence value (a.u.) compared to the fluorescence signal of the diploid cells in G1 was used to assign a ploidy of 3n. An increase of 100% of the fluorescence value (a.u.) compared to the fluorescence signal of the diploid cells was used to assign a ploidy of 4n.

### Long read sequencing and HGT identification

Yeast cells were grown overnight in liquid YPD medium. Genomic DNA was extracted using Qiagen Genomic-Tips 100/G according to the manufacturer’s instructions. The MINION sequencing library was prepared using the SQK-LSK108 sequencing kit according to the manufacturer’s protocol. The library was loaded onto a FLO-MIN106 flow cell and sequencing was run for 72 hours. We performed long read basecalling and scaffolding using the pipeline LRSDAY^40^. The Canu assembler mostly merged the different haplotypes and thus prevented to produce long-read phased genomes. The dotplots were generated by using mummerplot^41^. The annotated non-reference regions on chromosome XIII of OS1364 were extracted from the fasta file of the collapsed assembly genomes (the haplotypes were not phased) and blasted by using the application “blastn” at https://www.ncbi.nlm.nih.gov/ against the database of ascomycetes.

### Sporulation monitoring and spore viability

Yeast strains were patched on YPD plates and incubated at 30 °C for 24 hours. From the patch a streak for single was done and incubated at 30 °C for 48 hours, then single colonies were isolated and grown overnight at 30 °C in 10 mL of liquid YPD in a shaking incubator at 220 rpm. The following day, single colonies were inoculated in different tubes containing 10 mL of the pre-sporulation medium SPS (1% peptone, 1 % potassium acetate, 0.5% yeast extract, 0.17% yeast nitrogen base, 0.5% ammonium sulfate, 1.02% potassium biphthalate) and kept at 30°C in a shaking incubator for 24 hours. Tubes were then centrifuged and washed three times with sterile water and cells were resuspended in erlenmeyer flasks containing 25 mL of KAc 2% to reach a final OD of 1, and incubated at 23°C in a shaking incubator. Sporulation was monitored by DAPI staining of the sporulation cultures to detect cells which have passed the first meiotic division, and contain two nuclei, or the second meiotic division, and contain four nuclei. The cells were first washed with water and then fixed with EtOH 70 % overnight. Following, the cells were stained with DAPI and 10 μL of stained cells were spread on a microscope slide and incubated in the dark for 40 minutes. To infer the efficiency of meiotic progression the cells were analysed by fluorescence microscopy and scored as having one, two or four nuclei, which indicates if they have not progressed after MI (one nucleus) or have progressed after MI (two nuclei) or MII (four nuclei). For assesing the spore viability, the spores were collected from the sporulated cultures and incubated between 30-60 minutes in 100 μL of zymolase solution in order to perform spore dissection. At least 400 spores per sample were dissected on YPD plates. Spore viability was assessed as the number of spores forming visible colonies after 4 days of incubation at 30 °C.

### Plasmid engineering and genome editing with CRISPR/Cas9

The multi-deletions of *URA3*, *NDT80* and *SPO11* were engineered by using CRISPR/Cas9 genome editing. The plasmid harbouring Cas9 was obtained from Addgene pUDP004^42^ and linearized with BsaI. The resistance to acetamide was replaced with the resistance cassette to Kanamycin. The gRNA with the necessary nucleotide for self-cleavage was designed on UGENE^43^ and ordered as a synthetic oligo from Eurofins Genomics (^™^). The synthetic oligo was cloned into the plasmid backbone by using the Gibson assembly kit (NEB, Gibson Assembly^®^) and the ligation reaction was carried out for 1 hour at 50 °C. The assembled plasmid was transformed into DH5-alpha competent bacteria by heat shock and the bacteria were incubated in 3 mL of LB broth for 1 hour to induce the synthesis of the antibiotic resistance molecules and then plated on LB plates containing 100 μg/μL of ampicillin. The following day, cells were screened by polymerase chain reaction (PCR) using primers to validate the correct golden gate assembly of the construct. Successfully transformed bacterial colonies were inoculated in LB broth containing 100 μg/uL of ampicillin and incubated overnight at 37 °C. Cells were harvested from the overnight incubation and the plasmid was extracted using the QIAprep Spin Miniprep Kit following the manufacturer’s instructions.

The 120 bp cassettes used for the deletion of *URA3*, *NDT80* and *SPO11* were designed to be flanking 60 bp upstream and downstream the candidate gene which harbours the cutting site for Cas9 and ordered as a unique synthetic oligo at Eurofins Genomics. The forward and reverse cassettes were mixed at equimolar ratio, heated at 95 °C for 15 minutes and then cooled down at room temperature to be ready for the transformation. Yeast samples were transformed using between 1-15 ug of cassette, at least 200 ng of CRISPR/Cas9 plasmid and following the protocol from^42^. Cells were then plated on selective media containing kanamycin (400 ug/mL) and incubated at 30 °C for 3 - 7 days. Candidate transformed clones were validated by PCR using primers designed on the outside regions of the deleted genes. Positive clones were streaked for single on YPD and grown for 2 days at 30 °C to allow plasmid loss. Plasmid loss was confirmed by plating again the colonies in the selective medium and positive ones were patched on YPD and stored at −80 °C in 25% glycerol tubes.

### RTG selection by *URA3*-loss assay

A *URA3* cassette was introduced in one *LYS2* locus on chromosome II in *URA3* knock-out OS1364 and OS1431 strains by using the classical lithium acetate protocol. The correct insertion was validated by checking for restored prototrophy on synthetic media lacking uracil and by PCR. Positive clones were stored at −80 °C in 25% glycerol tubes. For performing the *URA3*-loss assay, cells from the frozen stocks were patched on YPD plates and incubated at 30 °C for 2 days. Cells were then streaked on plates lacking uracil and incubated at 30 °C for 2 days. Single colonies were picked and sporulation was induced following the protocol described in the paragraph “Sporulation monitoring and spore viability”. Cells from the sporulation cultures were taken at 0, 6 and 14 hours after sporulation induction, washed three times with YPD and incubated in YPD for 12 hours at 30 °C without shaking. Dilutions of the YPD liquid culture were spotted onto YPD plates and an appropriate dilution was chosen for each strain and plated on 5-FOA (2% dextrose, 0.675% yeast nitrogen base, 0.088% uracil drop-out, 0.005% uracil, 2 % agar, 0.1% 5-FOA plates. The plates were incubated at 30 °C for 2 days and colonies growing on 5-FOA plates were counted at all the time points. We calculated the LOH rate at the three time points (T0, T6 and T14) as the ratio between the % of cells growing on 5-FOA and the respective percentage of cells growing on YPD, according to the following equations:

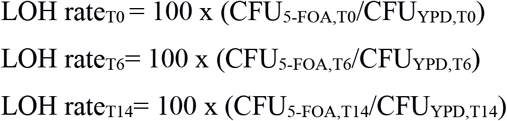

where CFU_5-FOA,T0_ is the number of colony forming units on 5-FOA at T0 and CFU_YPD,T0_ is the number of colony forming units on YPD at T0. These LOH rates were used to calculate: 1) the fold-increase of cells experiencing LOH by dividing the LOH rate at T6 or T14 by the LOH rate at T0 (LOH ratio); 2) the absolute difference of LOH by subtracting the LOH rate at T0 to the respective LOH rates at T6 and T14.

### RTG selection by natural phenotypes

Sporulation of wild-type hybrids was induced as described in “Sporulation monitoring and spore viability” for a time window compatible with RTG (14 hours). Cells were withdrawn from the sporulation medium before the appearance of MI cells to avoid plating committed meiotic cells. At the appropriate time point, cells were shifted from sporulation medium to liquid YPD. Part of the cells were incubated between 2 - 3 hours to induce RTG but not budding and separation of the mother-daughter RTG pairs, while others were incubated longer to allow complete budding and separation of candidates M-D RTG pairs. Then, the samples were plated on modified YPD media (YPD 0.5: 1% yeast extract, 2% peptone, 0.5% dextrose. YPD 1: 1% yeast extract, 2% peptone, 1% dextrose) and plates were incubated at 30 °C and monitored daily for colony formation and morphology variation. Mother-daughter RTG pairs were selected as single colonies displaying two sectors with different morphology, of which one resembled the wild-type phenotype whereas the other was divergent. Cells from each side of the sectors were taken with a wooden stick and streaked on their respective modified YPD to limit contamination from cells of the other sector, and incubated at 30 °C for 2 days. Then, a single colony was taken, patched on YPD and incubated at 30 °C for 2 days and finally stored in 25% glycerol tubes at −80 °C. Pictures of the sectoring colonies on the plates were taken with a stereomicroscope Discovery v.8 Zeiss.

### CNVs detection in parental strains and RTG samples

Short reads Illumina sequencings of were performed at the genomic platform of the Institute Curie. Reads were mapped to the S288C reference genome with the bwa-mem algorithm. Optical duplicates of the sequencing were removed using “samtools rmdup”. Processed BAM files were then indexed using “samtools index” and coverage was extracted using “samtools depth”. Coverage along the chromosomes was plotted using in-house R scripts in which sliding windows of non-overlapping 10 kbp were used to calculate a local average coverage, which was then normalized with the median coverage along the chromosome. Genome-wide coverage profiles were manually inspected to detect artifacts due to “smiley pattern”^5^ or lower coverage affecting only small chromosomes. Aneuploidies and large CNVs were detected by manually inspecting the log2 coverage profiles, while shorter CNVs (< 1 Kbp) were detected by computing the median coverage across sliding windows of non-overlapping 1 Kbp. Regions showing a coverage variation with respect to the median of their respective chromosome were then matched with the previously detected CNVs.

### LOH detection in RTG samples

To identify LOHs in the evolved RTGs, we first generated a list of reliable markers represented by all the heterozygous positions between the reference S288C strain and the two polyploid parental genomes (OS1364 and OS1431). Reads mapping and post-processing of sequenced parental strains, T0 and RTG samples were performed as described in the previous paragraph “CNVs detection in parental strains and RTG samples”.Variant calling in the parental strains was performed by using Freebayes (v1.3.1-19) with the options “-p” to set the appropriate ploidy. The parental vcf files were then filtered to include only “SNP” markers with quality >20 and depth >10 using bcftools with the options “TYPE=snp, QUAL > 20, DP > 10”. Variant calling on the evolved RTG and control (T0) clones was done using Freebayes with the options “-p” to set the appropriate ploidy, “-@” to call variants at the previously identified heterozygous positions and the additional options “-m 30, -q 20 -i -X -u” for minimum depth, quality and to exclude complex variants. Vcfs were then filtered with bcftools to take only “SNP” variants. As an additional filtering step, the parental and samples vcf files were intersected using bedtools “-intersect” and variants with shared positions were extracted from the vcf of the RTG evolved samples. Next, we used an in-house R script to compare the frequencies of the alternative and reference alleles of each marker in the evolved samples and in the parental ones. We identified putative recombined regions as the ones showing an allele frequency shift in the markers, independently of its direction. Heterozygous markers present in the parental strain but not in the RTG samples were considered to have reached a fixed reference homozygous genotype if the position was covered during sequencing. Moreover, the allele of the marker tagged as being in an LOH region was compared with the known allele of the marker present in the ancestral strain and those not matching the expected allele were filtered out. Given the complexity of polyploids analysis, we decided to use a stringent threshold and we annotated regions as LOH only if they contained at least 9 consecutive recombined markers regardless of the directionality of the allele that is gain or lost due to recombination. In this way, we avoided calling false positive LOHs at the cost of decreasing our resolution. Among those LOHs, the ones overlapping for >80% of their length among >=70% of the samples considering each datasets separately (wild-type RTG, *LYS2/URA3* selected RTG and *ndt80Δ* RTG) were filtered out. Additionally, LOH overlapping for >=90% of their length with the ones found in the T0 sampleswere also excluded. LOHs spanning the *LYS2/URA3* locus on chromosome II were not excluded but that chromosome was not counted for calculating the percentage of markers lying in LOH regions. Moreover, LOHs already present in the parental samples engineered with the *LYS2/URA3* system, or in which *NDT80* was deleted, were removed from the respective derived samples. LOHs in subtelomeric regions were excluded due to the unreliability of mapping in these highly repetitive regions. Finally, the list of annotated CNVs in core parts of the genome of the parental strains was used to filter the LOH detected. The plots of the LOH distribution were done using an in-house R script implemented with ggplot2 (v3.6.1) that takes into account the genotype shift of each marker in the LOH blocks.

### Inferring the parental haplotypes from the mother-daughter RTG pairs

One M-D RTG pair derived from OS1364 was used as a proof of concept to phase a region on the left arm of chromosome IX where recombination likely resulted from a cross-over between two homologs. The genotype of each heterozygous marker was inferred based on the allele frequency (AF) shift in the M-D pair. For example, when we detected an AF shift toward the alternative allele in a marker, that is, the RTG sample has gained one alternative allele, the genotype of the allele was assigned as the reference for that marker. We followed the same approach when we detected an AF shift toward the reference allele and assigned in this case the “alternative” genotype. Then, the information of these recombined haplotypes was used to infer the third haplotype based on the allele dosage, which is the number of “Ref” or “Alt” copies estimated for each heterozygous marker. For instance, if the initial genotype was “Ref/Alt/Alt” and the AF shift detected was toward the reference allele, the third unknown haplotype harboured an “Alt” allele and the two recombining haplotypes had respectively a “Reference” and “Alternative” allele. The information of the recombined haplotypes were validated on a second RTG M-D pair having a recombination event spanning the same region.

### Microplate cultivations

The osmotic stress and ethanol tolerance were assessed with microcultures in media containing 25% (w/v) sorbitol and 8% (v/v) ethanol, respectively. The microcultures were carried out in 100-well honeycomb microtiter plates at 25 °C (with continuous shaking), and their growth dynamics were monitored with a Bioscreen C MBR incubator and plate reader (Oy Growth Curves Ab, Finland). The wells of the microtiter plates were filled with 300 μL of YPM medium (1% yeast extract, 2% peptone, 1% maltose) supplemented with sorbitol (25%) and ethanol (8% v/v). Control cultivations in media without sorbitol or ethanol were also carried out. Precultures of the strains were started in 20 mL YPM medium and incubated at 25 °C with shaking at 120 rpm overnight. We measured the optical density at 600 nm, and pre-cultures were diluted to a final OD_600_ value of 3. The microcultures were started by inoculating the microtiter plates with 10 μL of cell suspension per well (for an initial OD_600_ value of 0.1) and placing the plates in the Bioscreen C MBR. The optical density of the microcultures at 600 nm was automatically read every 30 min. Four replicates were performed for each strain in each medium. Growth curves for the microcultures were modelled based on the OD_600_ values over time using the ‘GrowthCurver’-package for R^44^.

### Flask-scale very high gravity wort fermentations

50 mL-scale fermentations were carried out in 100 mL Schott bottles capped with glycerol-filled airlocks. Yeast strains were grown overnight in 25 mL YPM medium at 25 °C. The pre-cultured yeast was then inoculated into 50 mL of 32 °P wort made from malt extract (Senson Oy, Finland) at a rate of 7.5 g fresh yeast L^-1^. Fermentations were carried out in duplicate at 20 °C for 13 days. Fermentations were monitored by mass lost as CO_2_. The alcohol content of the final beer was measured with an Anton Paar density meter DMA 5000 M with Alcolyzer beer ME and pH ME modules (Anton Paar GmbH, Austria).

### 2-L scale high-gravity wort fermentations

Strains were characterized in fermentations performed in a 20 °P high gravity wort at 20 °C. Cultures were propagated essentially as described previously^45^, with the use of a “generation 0” fermentation prior to the actual experimental fermentations. The experimental fermentations were carried out in triplicate, in 3-liter cylindroconical stainless steel fermenting vessels, containing 2 liters of wort medium. The 20 °P wort (98 g of maltose, 34.7 g of maltotriose, 24 g of glucose, and 6.1 g of fructose per liter) was produced at the VTT Pilot Brewery from barley malt and malt extract (Senson Oy, Finland). Fermentations were inoculated at a rate of 5 g fresh yeast L^-1^ (corresponding to approximately 15 × 10^6^ viable cells · mL^-1^). The wort was oxygenated to 10 mg · L^-1^ prior to pitching (oxygen indicator model 26073 and sensor 21158; Orbisphere Laboratories, Switzerland). The fermentations were carried out at 20 °C until the alcohol level stabilized, or for a maximum of 15 days. Wort samples were drawn regularly from the fermentation vessels aseptically and placed directly on ice, after which the yeast was separated from the fermenting wort by centrifugation (9,000 × g, 10 min, 1 °C). Samples for yeast-derived flavour compound analysis were drawn from the beer when fermentations were ended. Cells viability was measured by propidium iodide staining of the cells that were collected at the end of the fermentations using a Nucleocounter ^®^ YC-100^™^ (ChemoMetec, Denmark).

### Chemical analysis of fermentable sugars

The concentrations of fermentable sugars (maltose and maltotriose) were measured by HPLC using a Waters 2695 separation module and Waters system interphase module liquid chromatograph coupled with a Waters 2414 differential refractometer (Waters Co., Milford, MA, USA). A Rezex RFQ-Fast Acid H^+^ (8%) LC column (100 × 7.8 mm; Phenomenex, USA) was equilibrated with 5 mM H2SO4 (Titrisol, Merck, Germany) in water at 80 °C, and samples were eluted with 5 mM H2SO4 in water at a 0.8 mL · min^-1^ flow rate. The alcohol level (% vol/vol) of samples was determined from the centrifuged and degassed fermentation samples using an Anton Paar density meter DMA 5000 M with Alcolyzer beer ME and pH ME modules (Anton Paar GmbH, Austria).

### Chemical analysis of aroma compounds in fermented beers

Yeast-derived higher alcohols and esters were determined by headspace gas chromatography with flame ionization detector (HS-GC-FID) analysis. Four-milliliter samples were filtered (0.45 μm pore size) and incubated at 60 °C for 30 min, and then 1 ml of gas phase was injected (split mode, 225 °C, split flow of 30 mL · min^-1^) into a gas chromatograph equipped with an FID detector and headspace autosampler (Agilent 7890 series; Palo Alto, CA, USA). Analytes were separated on a HP-5 capillary column (50 m by 320 μm by 1.05 μm column; Agilent, USA). The carrier gas was helium (constant flow of 1.4 mL · min^-1^). The temperature program was 50 °C for 3 min, 10 °C · min^-1^ to 100 °C, 5 °C · min^-1^ to 140 °C, 15 °C · min^-1^ to 260 °C, and then isothermal for 1 min. Compounds were identified by comparison with authentic standards and were quantified using standard curves. 1-Butanol was used as an internal standard.

## Acknowledgements

This work was supported by Agence Nationale de la Recherche (ANR-13-BSV6-0006-01, ANR-18-CE12-0004, ANR-15-IDEX-01), Fondation pour la Recherche Médicale (EQU202003010413), UCA AAP Start-up Deep tech, CEFIPRA, convention CIFRE 2016/0582 between Meiogenix and ANRT. We thank D’Angiolo M. and Adekunle D. for their critical reading of the manuscript.

## Author contributions

S.M., A.N., G.L., conceived the project, S.M., K.K., A.N., B.G., G.L. designed the experiments, S.M., K.K. performed the experiments and analyzed the data, S.M. and G.L. wrote the paper with input from K.K., B.G. and A.N.

## Competing interests

A.N. and G.L. have a patent application on “Yeast strains improvement method” using return-to-growth (US20150307868A1).

## Data and materials availability

The phenotype data are available within the supplementary tables. The short reads have been deposited on the SRA archive https://www.ncbi.nlm.nih.gov/sra with project number PRJNA770168.

