## Supplementary figures 1-9 for "Unlocking the functional potential of polyploid yeasts"

### Affiliations

### Notes

[#]Current address: Institute for Food Technology and Food Chemistry Department of Brewing and Beverage Technology, Technical University, 13353, Berlin, Germany

Correspondence and requests for materials should be addressed to:

S.M. and G.L..

### Content:

Supplementary Figures 1-9

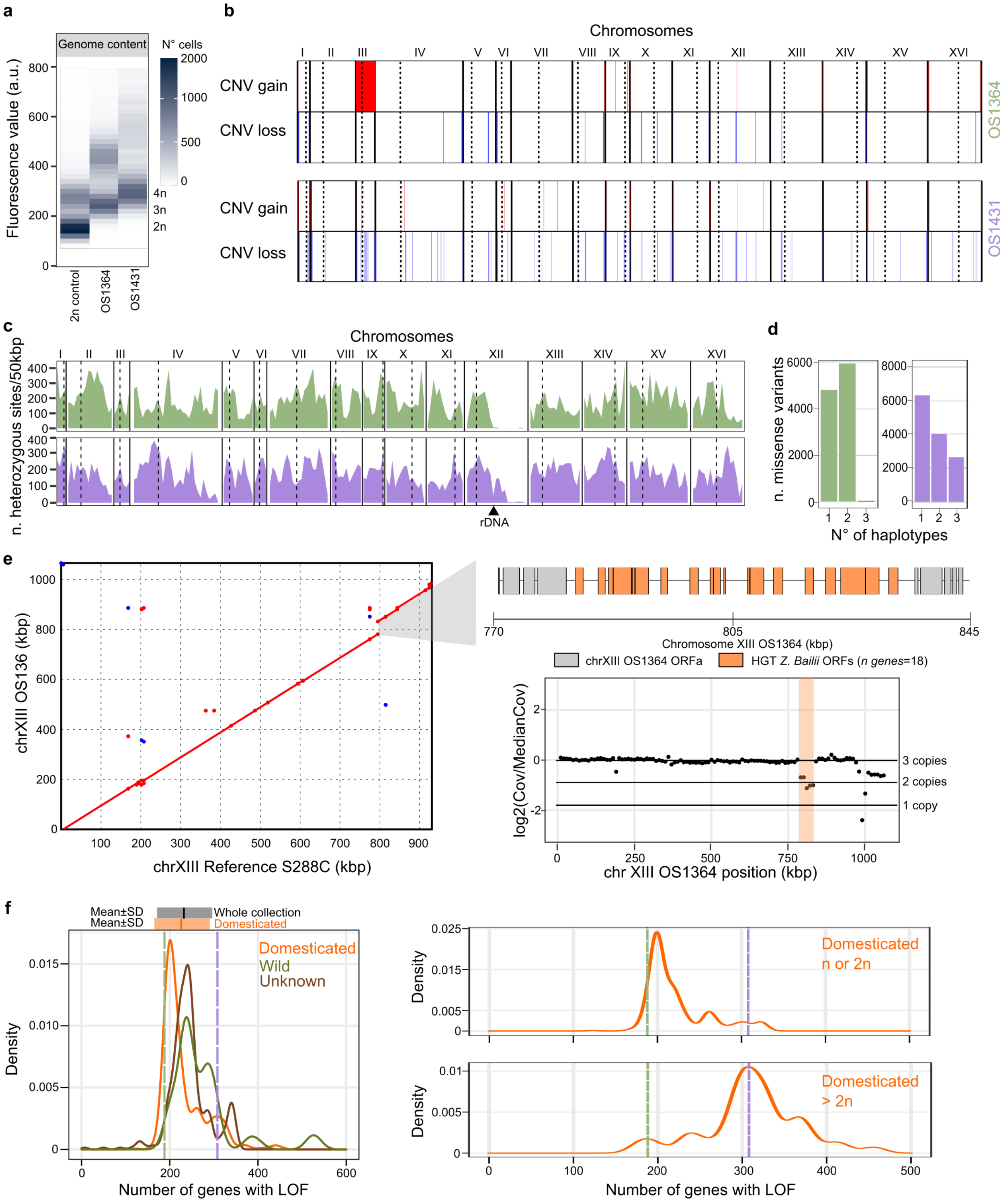


**Supplementary Figure 1. Genome characterization of OS1364 and OS1431 | (a)** Genome content estimated as the intensity of fluorescence after propidium iodide staining (y-axis) per each industrial hybrid as compared to a diploid control strain (2n). The labels on the right (3n-4n) are calculated based on the fluorescence of the 2n control strain. **(b)** Copy number variants (CNVs) detected in the two genomes. The aneuploidy on chromosome III of OS1364 is added as a whole large CNV. Each region with a copy number gain is highlighted in red and each region with a copy number loss is highlighted in blue **(c)** Genome-wide distribution of heterozygous markers calculated by using a non-overlapping window of 50 kbp. Dotted lines represent centromeres. **(d)** Barplots representing the number of missense variants (y-axis) based on the number of chromosomes on which they are found. Missense variants found on 3 haplotypes in OS1364 are related to chromosome III, which is present in 4 copies. **(e)** Left: dotplot of the alignment of OS1364 chromosome XIII (y-axis) against S288C chromosome XIII (x-axis). Right: annotation of the HGT region derived from

*Z. bailii* (top) and copy number of chromosome XIII with the HGT position highlighted in orange. **(f)** (Left) Distribution of the strains according to the respective number of genes with loss-of-function (LOF) (x-axis) across the 1011 *S. cerevisiae* collection. The distributions are colored according to the ecological origin of the strains. The mean (solid line) and the interval encompassing one standard deviation from the mean are reported on the top. (Right) As described for the plot on the left but using only domesticated strains divided by their genome content. Top: haploids or diploids. Bottom: higher ploidy. The dashed lines in all plots represent the number of genes with LOF in OS1364 (green) and in OS1431 (purple).

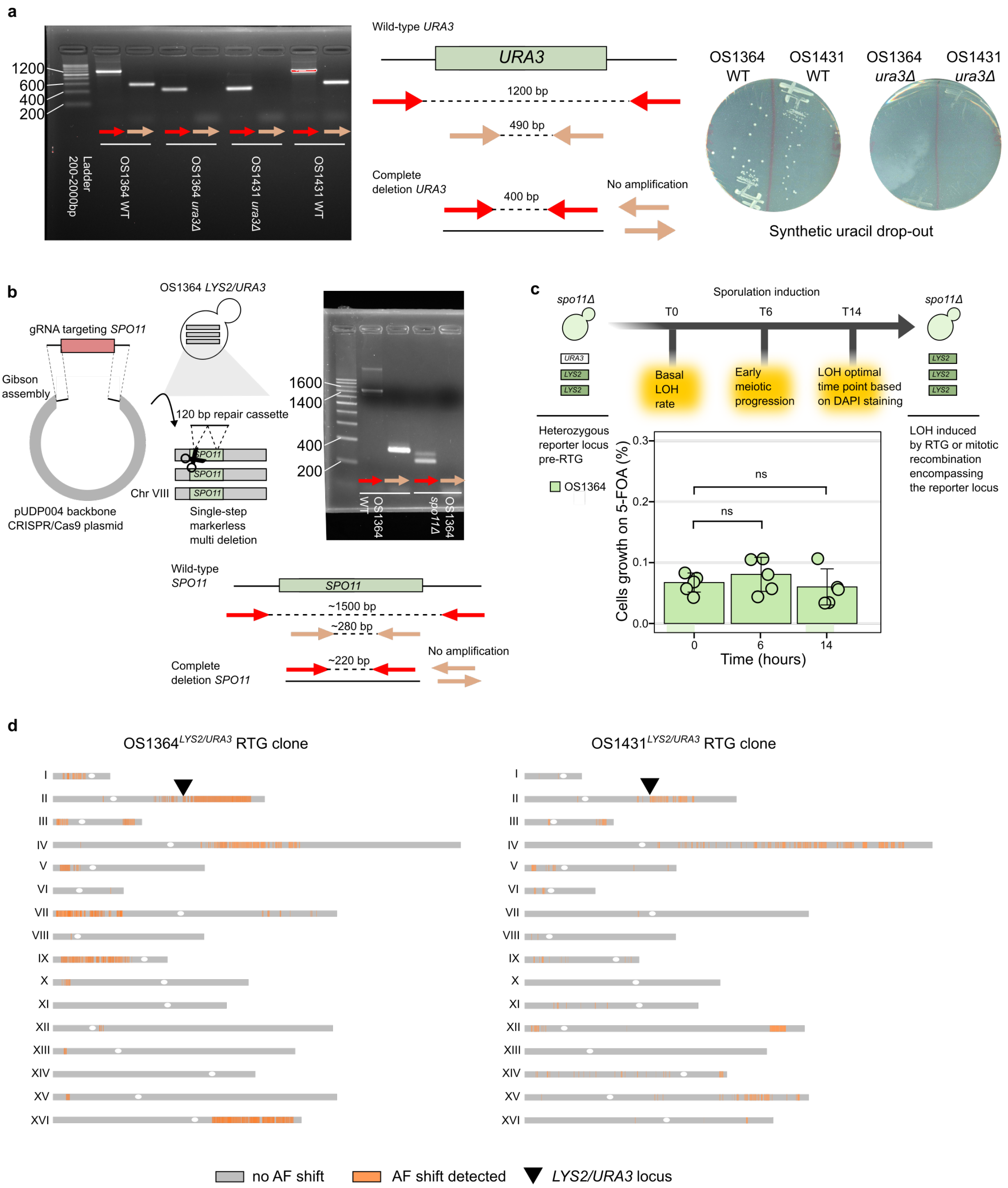


**Supplementary Figure 2. *URA3*-loss assay in polyploid hybrids | (a)** PCR validation of the simultaneous deletion of all the *URA3* genes in their native locus on chromosome V. An example is reported for both OS1364 and OS1431. The replacement of the genes by homologous-recombination with a repair cassette whose first and last 60 nucleotides are designed on the flanking regions of the gene produces a deletion which can be visualized by PCR as a band of reduced size, when using primers binding outside the gene (red), and by the absence of amplification when using internal primers (brown). **(b)** The same strategy was used for deleting all the gene copies of *SPO11* in OS1364 **(c)** *URA3*-loss assay performed with the mutant strain OS1364$^{LYS2/URA3,}$ *spo11Δ*. We did not detect increase in LOH rate neither at T6 (One-sided Wilcoxon rank-sum test, *p*-value > 0.05) nor at T14 (One-sided Wilcoxon rank-sum test, *p*-value > 0.05) as compared to the initial LOH rate (T0) **(d)** LOH map of two RTGs selected on 5-FOA in which identified LOH regions are highlighted in orange. Black triangles indicate the locus bearing the *URA3* selection marker. LOH regions are defined as tracts of heterozygous markers for which we detected an allele frequency (AF) shift as compared to their respective parental allele dosage (see methods).

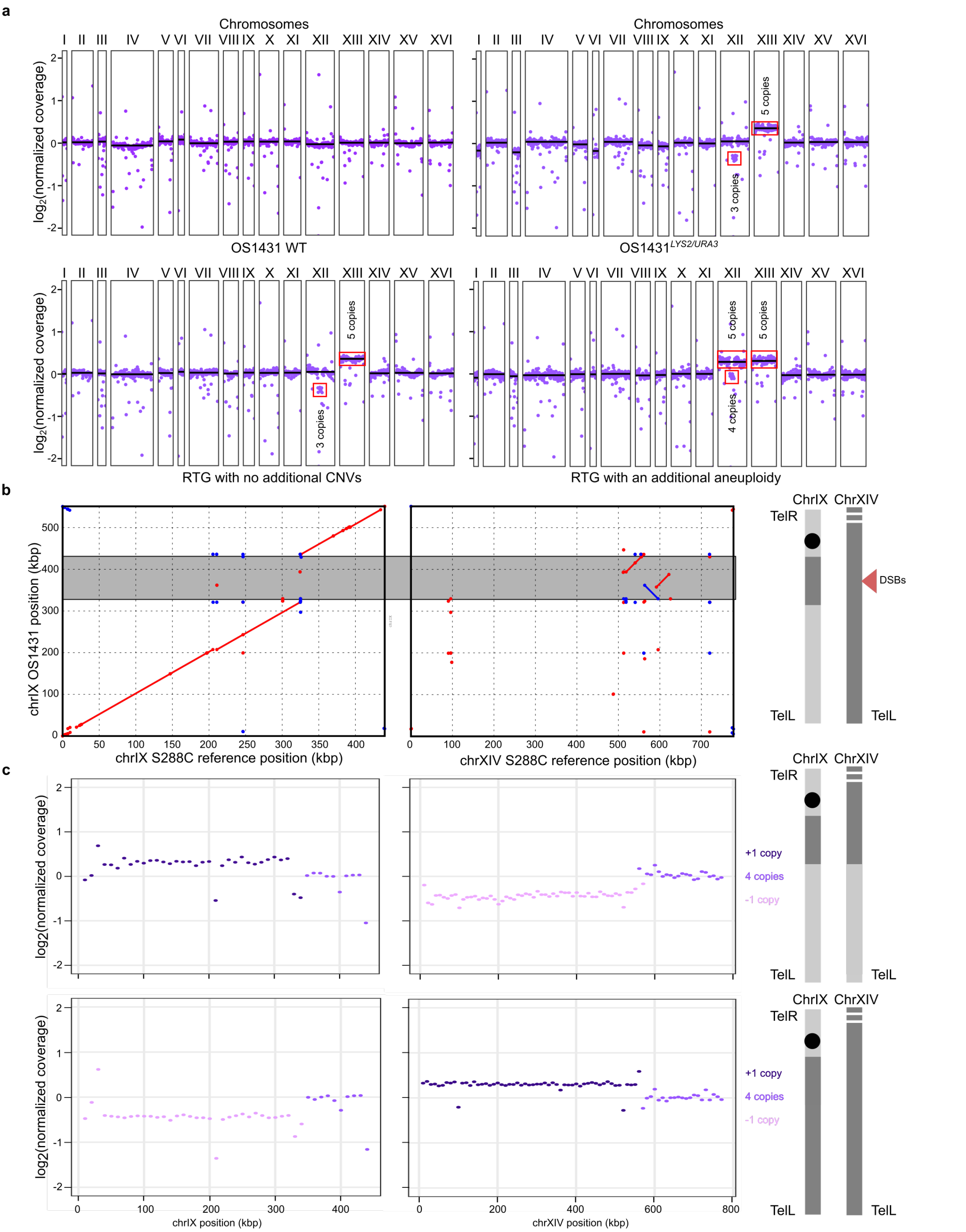


**Supplementary Figure 3. Genome content variation induced by CRISPR/Cas9 and RTG in OS1431$^{LYS2/URA3}$ | (a)** Genome-wide coverage plots of the parental OS1431, OS1431$^{LYS2/URA3}$ and its derived RTGs. The coverage profile of the CRISPR/Cas9 engineered OS1431$^{LYS2/URA3}$ and the parental OS1431 are reported on the top. Copy number variants and aneuploidies, which were not present in the parental OS1431 but were detected in the CRISPR/Cas9 engineered parent are highlighted. On the bottom: RTG-derived samples selected on 5-FOA without additional CNVs (left) or with an additional aneuploidy (right). The black lines represent the median coverage of each chromosome. **(b)** Dotplot highlighting a region of the parental OS1431 chromosome IX bearing a complex insertion from chromosome XIV (left). On the right: scheme illustrating how a DSB in the region of homology between the two chromosomes can give rise to the rearranged chromosomes reported below in **(c)**. **(c)** Each row represents the coverage profile of chromosome IX and chromosome XIV of one RTG sample bearing a CNV (left). On the right: scheme with the predicted chromosome structure generated by the RTG-induced recombination.

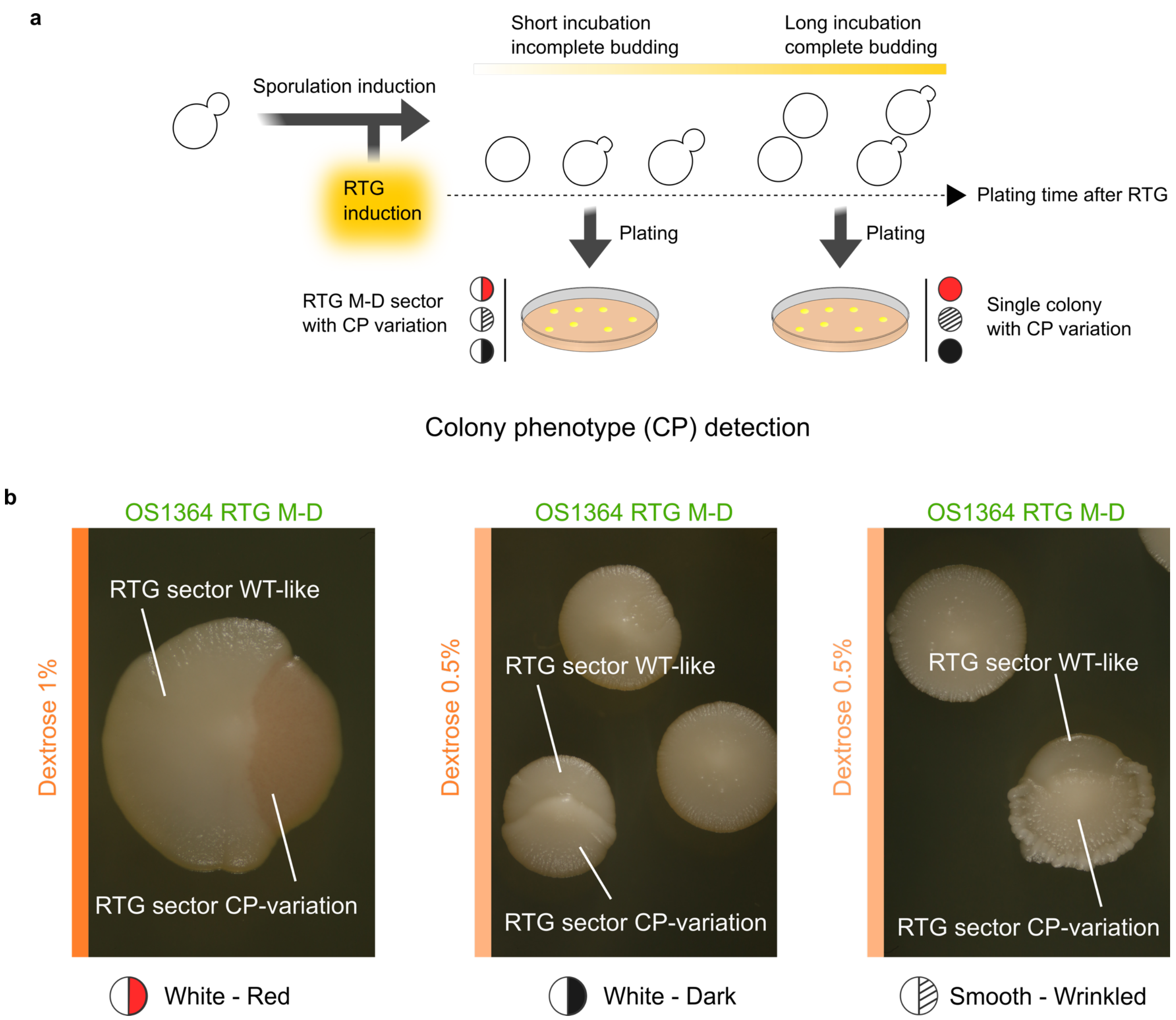


**Supplementary Figure 4. Identification of RTG Mother-Daughter (M-D) sector phenotypes | (a)** Different plating times allowed us to retrieve RTG sectors. If cells are kept in rich medium long enough as to induce RTG and complete budding, the M-D RTG cells will bud and will be found as single colonies on the plate. Viceversa, if cells are left in rich medium and plated before complete budding, RTG M-D can be retrieved as a colony with two sectors displaying different phenotypes. **(b)** Example of the three phenotypic classes of RTG M-D used for isolating RTG candidates in the parental OS1364 and in the OS1364$^{ndt80\Delta}$ genetic background.

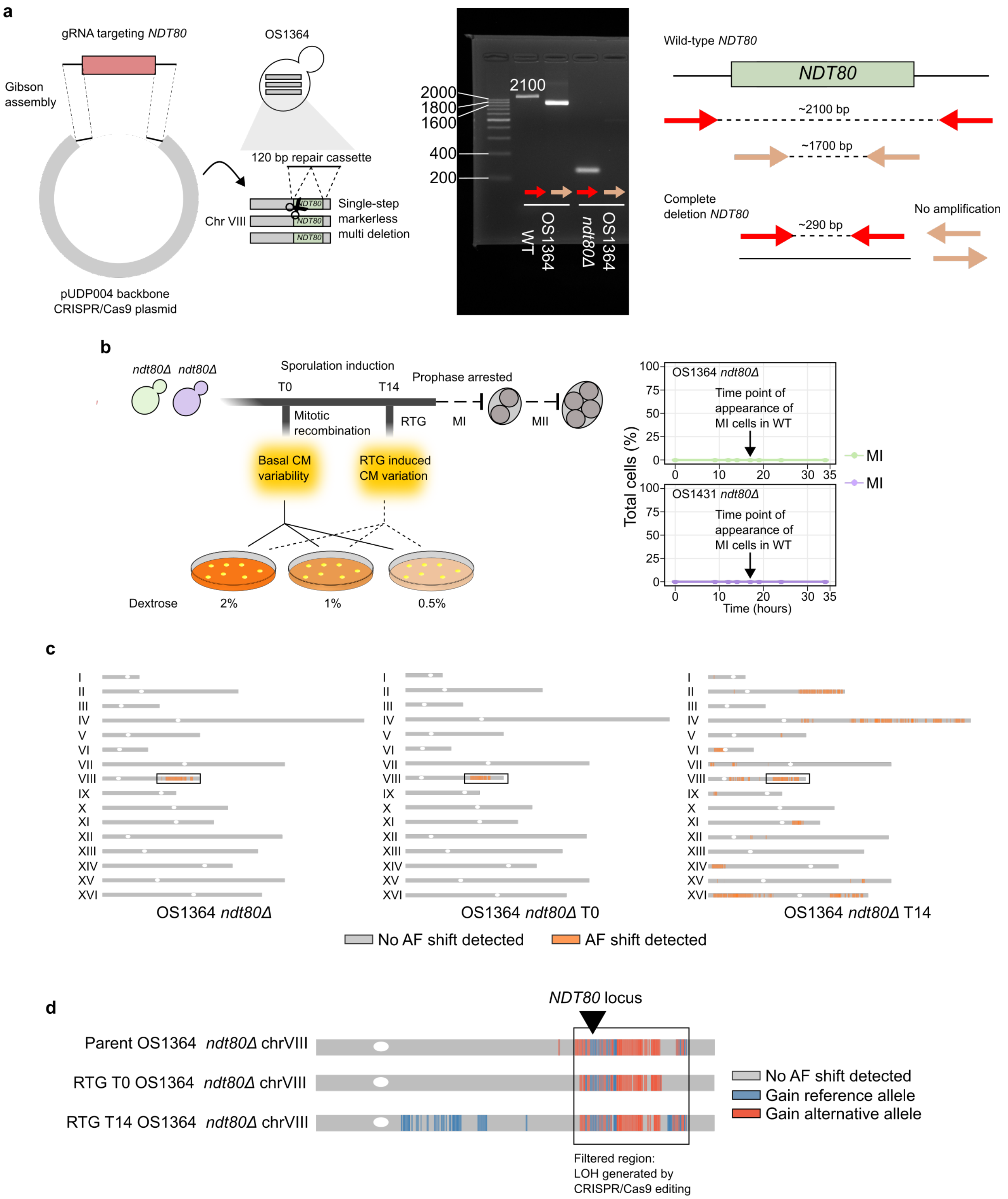


**Supplementary Figure 5. Genome engineering to generate *NDT80* mutants in OS1364 and OS1431 backgrounds | (a)** Deletion of *NDT80* using the same strategy reported in Supplementary Figure.2. A PCR validation example for OS1364 is reported, and it follows the same rationale used for *URA3* and *SPO11* deletions. **(b)** The *ndt80Δ* strains were evolved following the same RTG protocol used for the WT strains (left). Meiotic progression was monitored by DAPI staining and this approach confirmed the absence of cells passing the first meiotic division (MI) even at later time points (right). **(c)** Three maps of recombination events are reported for *ndt80Δ* RTGs of OS1364. The LOH encompassing the *ndt80* locus on chromosome VIII resulted from CRISPR/Cas9 engineering and is highlighted with a box. **(d)** Zoom-in of the recombination events on chromosome VIII in the samples reported above, which share a common LOH generated upon genome-editing.

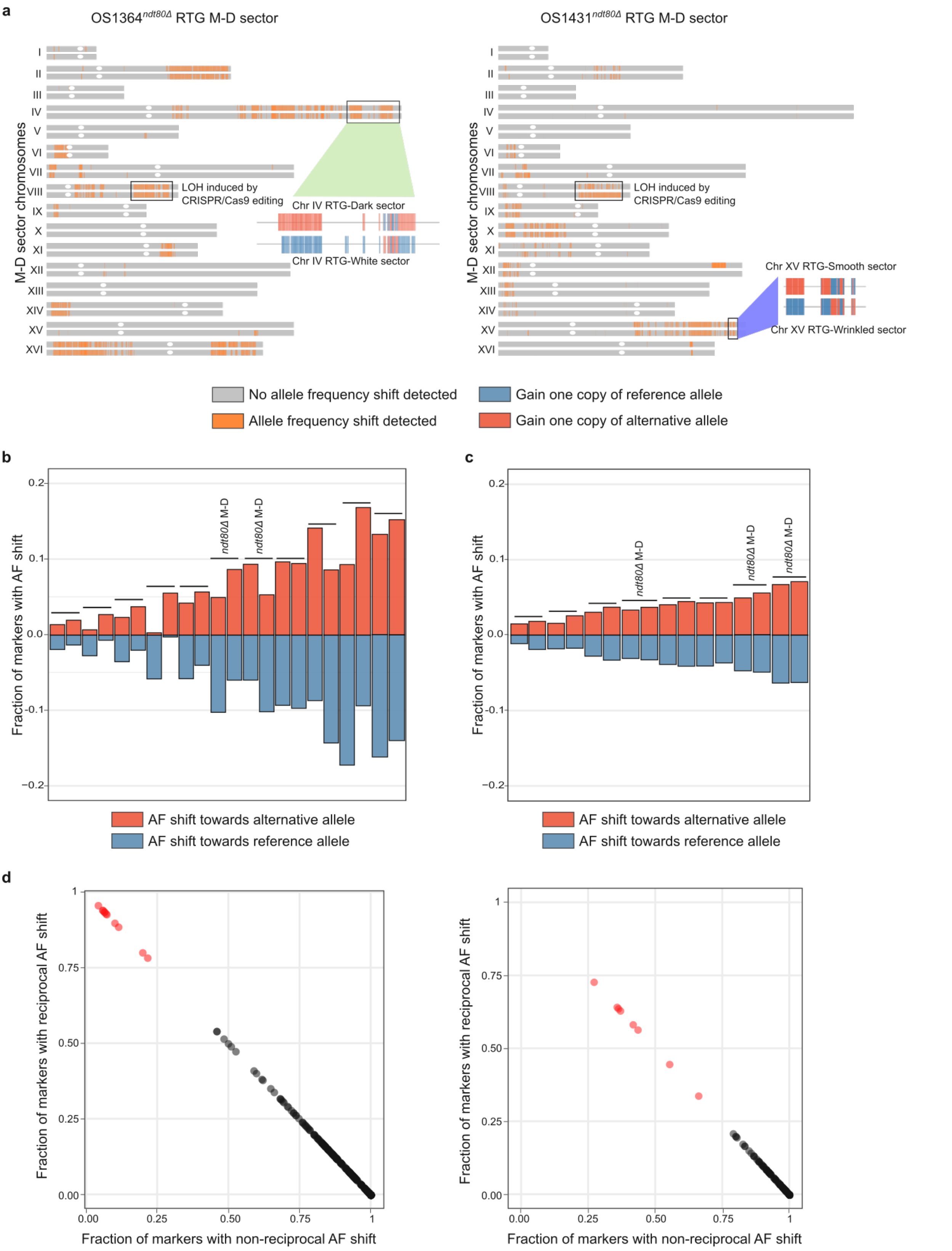


**Supplementary Figure 6. Recombination profile in RTG M-D sectors derived from WT and *ndt80Δ* strains | (a)** Recombination map of two M-D *ndt80Δ* RTG sectors derived from OS1364 and OS1431 with a zoom-in on two recombination events where the direction of allele frequency (AF) shift is reported for both RTG sectors. **(b)** Fraction of markers in which we detected an AF shift toward an increase of alternative allele (red) or parental allele (blue) as a result of recombination in OS1364 M-D RTG sectors. Each M-D RTG sector pair is reported sequentially from left to right and marked with a horizontal bar. Each bar represents one RTG sample of the pair. The region in which recombination was induced by CRISPR-Cas9 was removed from the count. **(c)** Fraction of markers in which we detected an AF shift toward an increase of alternative allele (red) or parental allele (blue) as a result of recombination in OS1431 M-D RTG sectors. Each M-D RTG sector pair is reported sequentially from left to right. Each bar represents one RTG sample of the pair. The region in which recombination was induced by CRISPR-Cas9 was removed from the count. **(d)** Scatter plot in which each point represents a mother-daughter RTG combination of the RTG library taking together the M-D RTG pairs derived from the WT and the *ndt80Δ* strains of OS1364 (left) and OS1431 (right). For each combination we calculated the fraction of heterozygous markers that have reciprocal (y-axis) or non reciprocal (x-axis) AF-shift. In red are represented the real RTG M-D pairs derived from the same colony while in black are represented all the other pair-wise combinations.

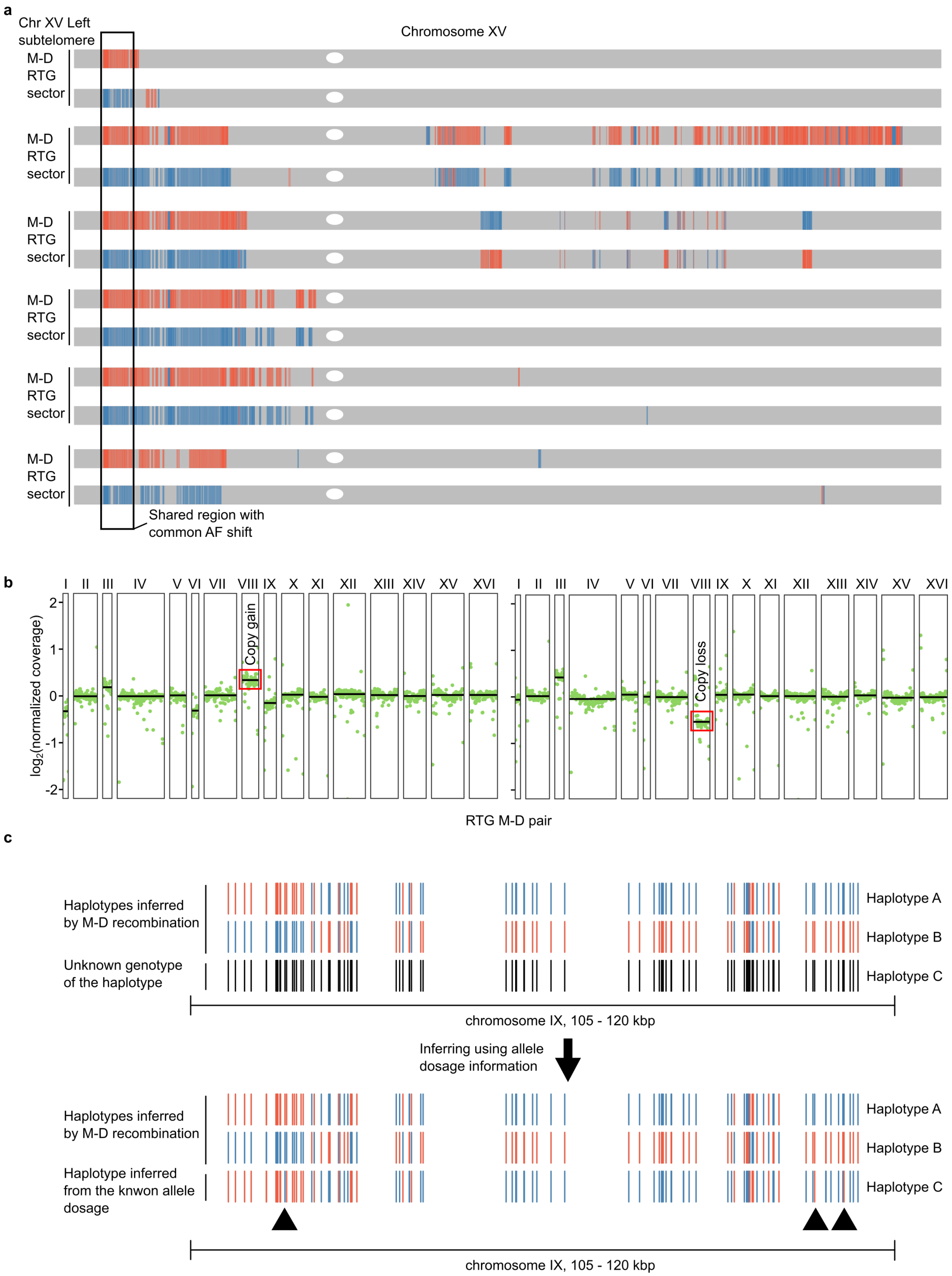


**Supplementary Figure 7. Analysis of recombinations in RTG M-D sectors | (a)** Recombination pattern in chromosome XV where the colors represent a shift of the genotype of the heterozygous marker toward the reference (blue) or alternative (red) allele. The rectangle highlights a shared region of recombination in six distinct M-D pairs with the same sectoring phenotype (dark-white). **(b)** Genome-wide coverage profiles where complementary gain (left)

and loss (right) of chromosome VIII in a OS1364 M-D pair is highlighted. Each plot represents one of the two samples. The black lines represent the median coverage of each chromosome. **(c)** Example of local haplotype reconstruction based on the recombination profile of chromosome IX of a M-D pair (top), which is used to infer the third haplotype by using the information of allele frequency for each heterozygous marker (bottom). Red represents gain of an alternative allele whereas blue represents gain of a reference allele. The black triangles point to heterozygous markers that are discordant between haplotype A and haplotype C.

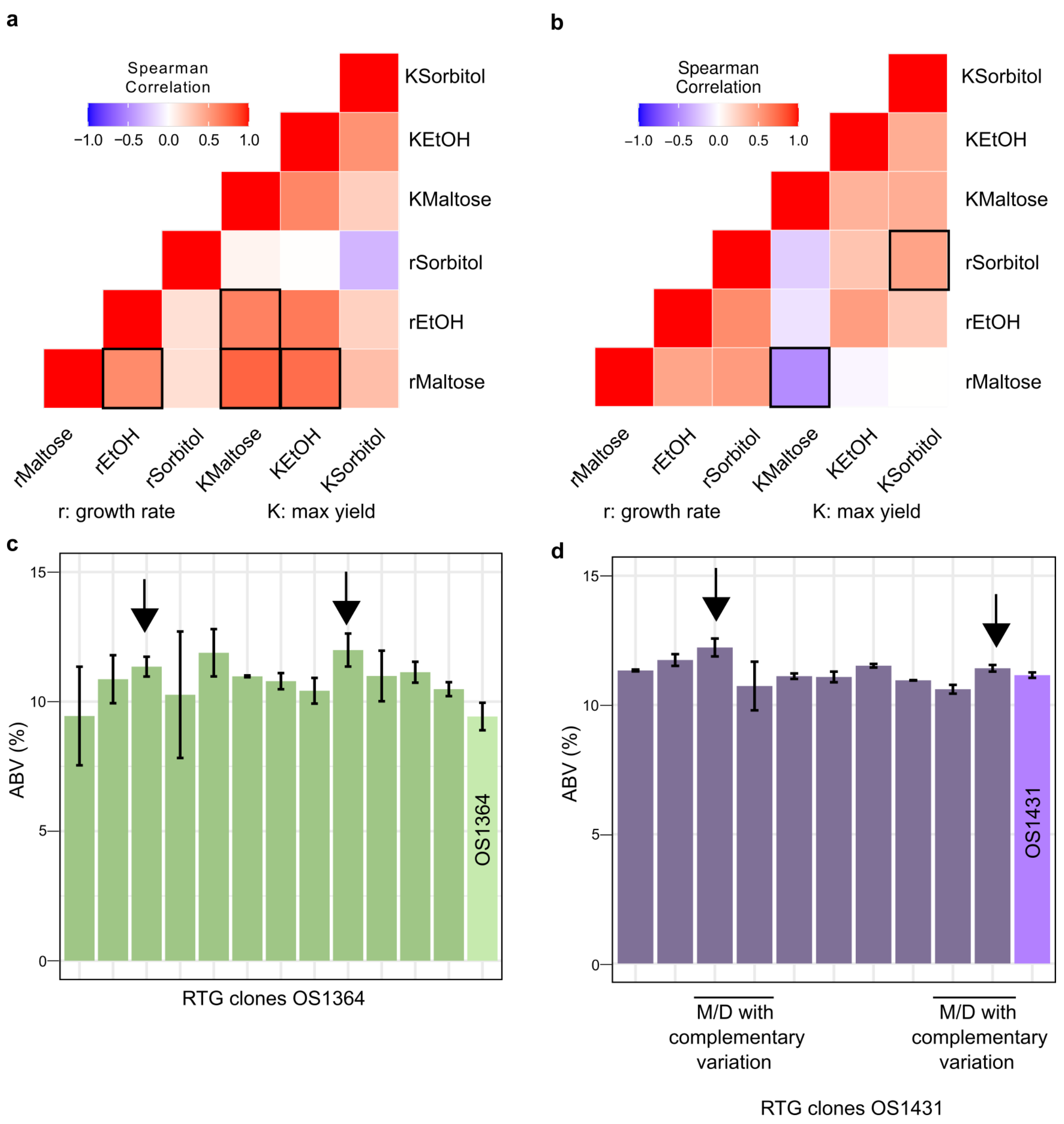


**Supplementary Figure 8. Phenotypic correlation and diversification analyses | (a)** Spearman correlation between phenotypic performances in stressor or maltose conditions measured in RTGs derived from the WT OS1364. Black squares denote significant associations calculated by using Bonferroni correction for multiple testing. **(b)** Spearman correlation between phenotypic performances in stressor or maltose conditions measured in RTGs derived from the WT OS1431. Black squares denote significant associations calculated by using Bonferroni correction for multiple testing. **(c)** Bar plots representing alcohol by volume (ABV%) produced at the end of the flask-scale fermentation experiment in OS1364 RTG clones. Black arrows point to the samples selected for the 2L scale fermentation. The error bar represents the standard deviation and the bar plot the mean of two indipendent fermentations. **(d)** Bar plots representing alcohol by volume (ABV%) produced at the end of the flask-scale fermentation experiment in OS1431 RTG clones. Black arrows point to the samples selected for the 2L scale fermentation. The error bar represents the standard deviation and the bar plot the mean of two independent fermentations.

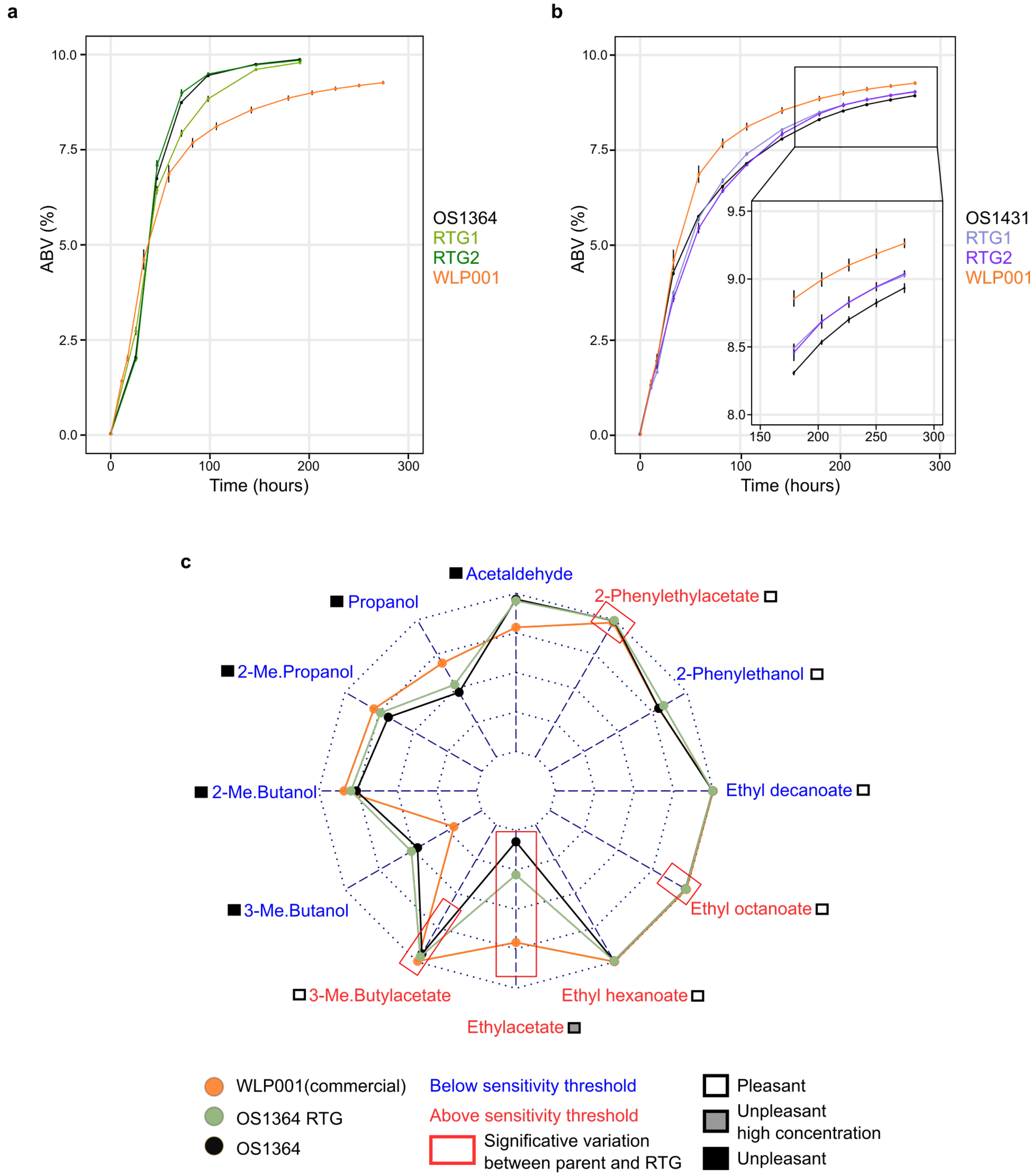


**Supplementary Figure 9. 2L fermentation kinetics and sensory quality of the end product | (a)** Fermentation profile of two RTGs (RTG1 = sm244, RTG2 = sm338) derived from OS1364. The alcohol by volume (ABV%) measured at different time points is reported on the y axis. WLP001 is the reference commercial strain. All the measures are an average of 3 technical replicates. **(b)** Fermentation profile of two RTGs (RTG1 = sm399, RTG2 = sm408) derived from OS1431. The alcohol by volume (ABV%) measured at different time points is reported on the y axis. WLP001 is the reference commercial strain. **(c)** Web chart representing the aroma profile of the beer fermented by sm338. Significant variations that cannot be spotted by eye are due to low sensory threshold (mg/L) compared to other flavours. The data relative to them are reported in **Supplementary Table 11**. Aromas below (blue) or above (red) sensory threshold are color-coded. The red squares highlight significant differences which are above the sensory threshold.
